# Insights from Fisher’s geometric model on the likelihood of speciation under different histories of environmental change

**DOI:** 10.1101/596866

**Authors:** Ryo Yamaguchi, Sarah P. Otto

## Abstract

The formation of new species via the accumulation of incompatible genetic changes is thought to result either from ecologically-based divergent natural selection or the order by which mutations happen to arise, leading to different evolutionary trajectories even under similar selection pressures. There is growing evidence in support of both ecological speciation and mutation-order speciation, but how different environmental scenarios affect the rate of species formation remains underexplored. We use a simple model of optimizing selection on multiple traits (“Fisher’s geometric model”) to determine the conditions that generate genetic incompatibilities in a changing environment. We find that incompatibilities are likely to accumulate in isolated populations adapting to different environments, consistent with ecological speciation. Incompatibilities also arise when isolated populations face a similar novel environment; these cases of mutation-order speciation are particularly likely when the environment changes rapidly and favors the accumulation of large-effect mutations. In addition, we find that homoploid hybrid speciation is likely to occur either when new environments arise in between the parental environments or when parental populations have accumulated large-effect mutations following a period of rapid adaptation. Our results indicate that periods of rapid environmental change are particularly conducive to speciation, especially mutation-order or hybrid speciation.

## Introduction

According to the ecological theory of speciation, populations diverge when exposed to different environmental conditions (Schluter 2000). Metaphorically, populations in different environments are thought to evolve phenotypically across different fitness landscapes, diverging over time as they approach different fitness peaks. This theory can be tested using reciprocal transplant experiments in which the performance of phenotypically divergent populations and their hybrid progeny are measured in both parental environments. The results of such studies often indicate that the environment matters, with phenotypic differentiation and the performance of hybrids depending on the environment (Table 5.1 in Schluter 2000).

An alternative theory of speciation – “mutation-order speciation” – proposes that populations diverging under similar selection pressures may ultimately become reproductively isolated because they accumulate different sets of mutations, some of which are incompatible with one another (Mani and Clarke 1990; Schluter 2009; Schluter and Conte 2009; Mendelson *et al.* 2014). The focus here is on the reduction in fitness caused by combining genes that have independently spread in different populations, so-called “Bateson-Dobzhansky-Muller” incompatibilities (BDMI), regardless of the environment in which hybrids are assayed.

Empirical studies have suggested that both divergent selection and parallel selection are relatively common (Table 5.2 in Schluter 2000), suggesting that both ecological and mutation-order speciation may underlie the formation of new species in nature. Ecological speciation is often inferred based on local adaptation of parental populations to different abiotic (e.g., soil chemistry) or biotic conditions (e.g., pollinator communities) (see Waser and Campbell 2004 for a review of evidence in plants). While such data provides evidence for ecology-dependent selection, it does not prove that ecological differences underly reproductive isolation, rather than intrinsic genetic incompatibilities (BDMIs). Additional evidence for ecological speciation comes from reciprocal transplant experiments where hybrids formed by backcrossing to the foreign parent are less fit than backcrosses to the local parent (Rundle and Whitlock 2001), as seen in sticklebacks (Rundle 2002). Alternative evidence for ecological speciation is provided when populations that have independently adapted to a novel environment and that are isolated from their ancestral populations remain interfertile with other such populations from similar environments (Schluter 2000). Such “parallel ecological speciation” has been found in a number of animal taxa, including sticklebacks and other fish, Phasmatodea walking-stick insects, and *Littorina* snails, but evidence in plants remains weak (Ostevik *et al.* 2012).

Empirical evidence for mutation-order speciation includes cases of reciprocal gene loss of duplicated genes in *Drosophila, Arabidopsis*, and yeast, where the loss of one gene copy or the other is thought to have occurred under similar selection pressures, reducing fitness in hybrids that lack the gene (see discussion and review of evidence by Muir and Hahn 2015). Genetic conflict, such as meiotic drive or cytonuclear conflicts, has also been shown to generate reproductive isolation in a manner consistent with mutation-order speciation in a wide variety of animal and plant species (Crespi and Nosil 2013).

Experimental evolution studies in the lab have also provided evidence for both ecological speciation and mutation-order speciation. In such studies, multiple independent cultures of organisms are allowed to evolve under either different environmental conditions or identical conditions, following which genetic incompatibilities are measured between the replicate populations. Evidence for ecological speciation has been obtained in studies that have adapted lines to different conditions. For example, Dettman *et al.* (2007) evolved multiple populations of the yeast *Saccharomyces cerevisiae* in either rich media with high salt or in minimal media with low glucose. In most replicates, hybrid diploids performed significantly worse than the strain that had adapted to a particular local environment. Similarly, Dettman *et al.* (2008) adapted *Neurospora crassa* to either high salinity or low temperature conditions, finding evidence for the first stages of parallel ecological speciation. When crossed, reproductive success was lower among hybrids formed from strains adapted to different environments than among hybrids from strains adapted to the current environment being tested.

Evidence for mutation-order speciation includes an experimental evolution study by Kvitek and Sherlock (2011) in *S. cerevisiae* involving multiple replicate populations propagated under glucose limitation. Genomic sequencing revealed that mutations in two genes (MTH1 and HXT6/HXT7) arose repeatedly in different lines but were never found together. Subsequent crosses established that adaptive mutations at each of these two genes were deleterious when combined, reducing the fitness of hybrids. Similarly, a study of *S. cerevisiae* adapting in the presence of the fungicide nystatin found that even the first-step mutations that arise during adaptation can be incompatible (Ono *et al.* 2017). When lines that had accumulated different resistance mutations were crossed, the double mutants performed less well than single mutants in many cases (“sign epistasis”). Sign epistasis was also observed in an experiment with the bacterium, *Methylobacterium extorquens*, evolving to tolerate a genetically-engineered change to how formaldehyde is oxidized (Chou *et al.* 2014). Again, adaptive mutations tended to work poorly when combined.

Previous theoretical studies have modeled the fitness of recombinant hybrids under divergent and parallel adaptation following a period of adaptation with *de novo* mutations (Barton 2001; Chevin *et al.* 2014; Fraïsse *et al.* 2016). These models have clarified how segregation of different adaptive alleles reduces hybrid fitness (“segregation load”), via the separation of coadapted alleles. Using Fisher’s geometric model to track fitness variability among hybrids between independently adapting populations, Chevin *et al.* (2014) found that segregation load (isolated from extrinsic, environment-dependent, fitness effects) is insensitive to whether adaptation occurred to the same or to different environmental optima (e.g., Figure 2 in Chevin *et al.* 2014). This work is a launching point for the current study, where we explore how the distribution of potential mutations and the speed of adaptation influence the mean fitness of hybrids, combining the segregation load and the mismatch to the environment in which hybrid and parental fitnesses are measured (i.e., both intrinsic and extrinsic incompatibilities; their Equation 1, see also Equation 3 of Barton 2001). Fraïsse *et al.* (2016) further found that Fisher’s geometric model can generate many of the empirical patterns of speciation studies, from Haldane’s rule to asymmetrical introgression.

**Figure 1:**
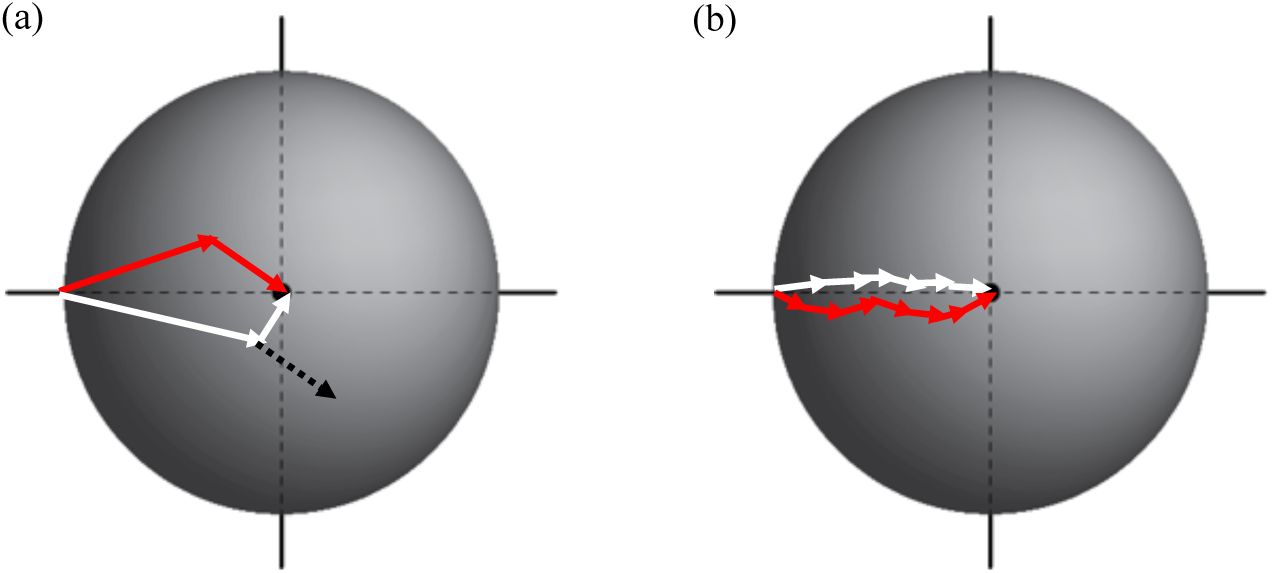
Conceptual figure illustrating why rapid environmental change is more likely to lead to reproductive isolation. Both panels show a two-dimensional phenotype with a new optimum at the center of the sphere, illustrating the adaptive paths taken by two populations (red and white). In (a), the environment shifted rapidly and favored large-effect mutations; in (b), the environment shifted slowly and favored small-effect mutations. Rapid environmental change (a) is more likely to generate genetic incompatibilities that reduce hybrid fitness because combining large-effect mutations can shift the offspring off of the fitness peak and because such mutations can have substantial negative pleiotropic effects, here measured as displacements along the y-axis (see dashed black arrow for one example hybrid combination). By contrast, slow environmental change (b) is more likely to lead to many small-effect mutations, which when combined create hybrids whose phenotypes remain near the peak.

**Figure 2.**
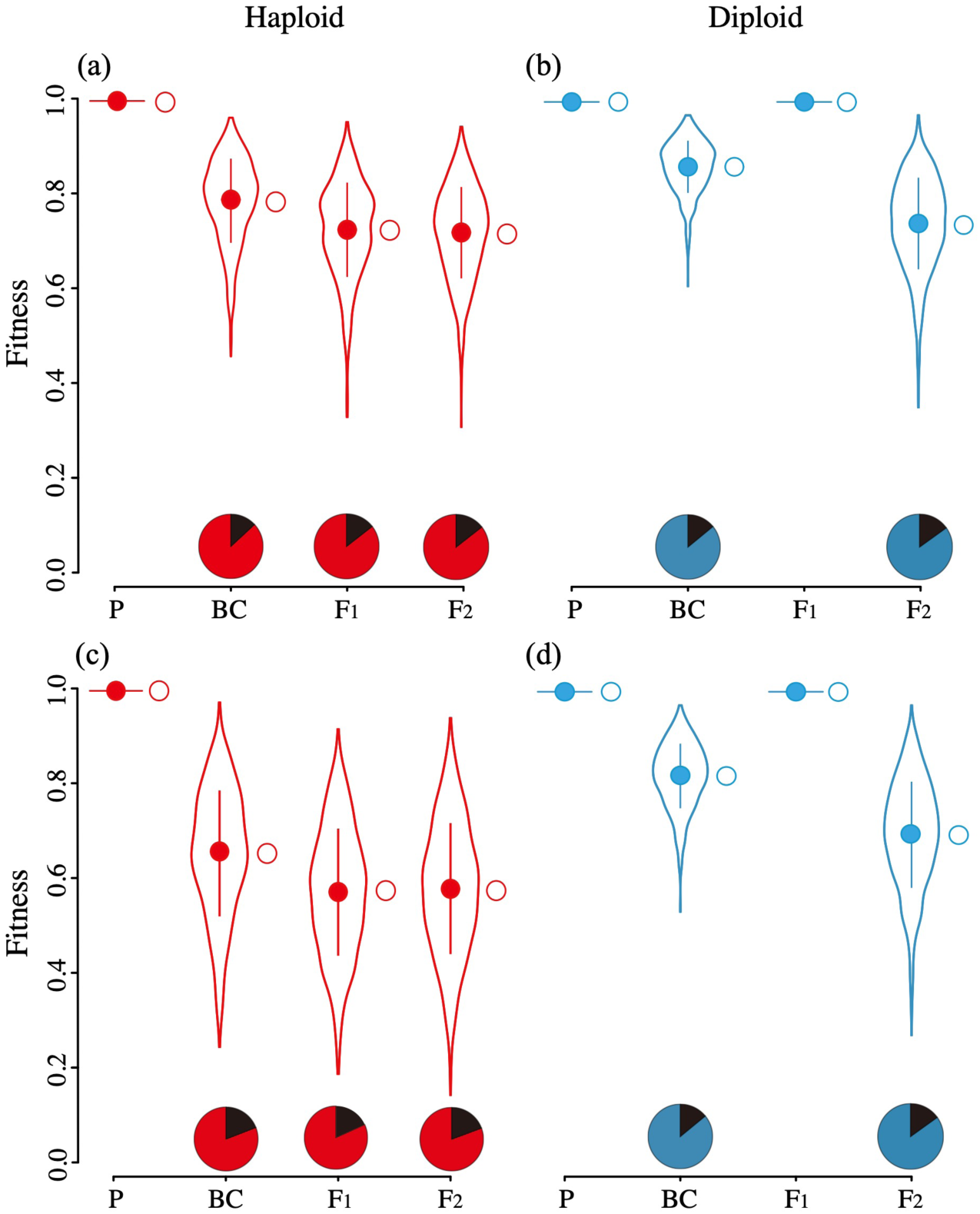
Mutation-order speciation and fitness of hybrids in (a, c) haploids and (b, d) diploids. We arbitrarily consider population 1 to be the “local” population used as the backcross parent, BC_L_ (backcross results using the “foreign” population 2, BC_F_, are similar). The phenotypic optimum was displaced from the origin along the x-axis by 2 units, after which the appearance and fixation of mutations was tracked for a total of (a) 360, (b) 388, (c) 144, (d) 246 substitutions (see Table S2). The majority of fitness loss accumulates during the first adaptation phase (open circles). Pie charts illustrate the fraction of the hybrid breakdown due to phenotypic change along the single axis that has undergone environmental change (black), while the remainder is due to departures from the optimum along other trait dimensions (due to pleiotropy, red in haploids, blue in diploids). Top panels show *λ*= 0.04 and bottom *λ*= 0.12. Parameters: *m*=1, *n*=10, *N*=2000, *q*=1.0, *k* = 2.0, with distributions and solid points representing the mean ± SD calculated from one simulation. 1000 individuals were generated for each hybrid type.

As noted in these two studies, large-effect mutations are more likely to contribute to genetic incompatibilities. This prediction is consistent with the observation that incompatibilities arise often in short-term experimental evolution studies, because those studies typically induce strong selection favoring large-effect mutations. Large-effect mutations are more likely to cause hybrid breakdown both because large-effect mutations can have large pleiotropic side effects and because combining such mutants can overshoot the optimal fitness (Figure 1). Our work thus aims to explore the environmental conditions that allow such strong genetic incompatibilities to arise, measuring how overall reproductive isolation, including both intrinsic and extrinsic isolation, depends on the speed and direction of environmental change, as well as the average effect size of new mutations.

In another recent study, Simon *et al.* (2018) looked not at the processes generating divergence between parental populations but at the fitness relationships between different hybrid genotypes. Given a set of divergent alleles between the parents, hybrid fitness depends on two main quantities, the hybrid index, *H* (the total proportion of alleles from one parent), and heterozygosity, *p*_12_ (the proportion of divergent sites that carry one allele from each parent population). Using a Brownian bridge approximation to walk from hybrids with only alleles from one parent (*H* = 0) to hybrids with only alleles from the other parent (*H* = 1) and using Fisher’s geometrical model to describe fitness, the authors predict the average fitness of all hybrid combinations (*H, p*_12_) given remarkably few summary statistics (the breakdown scores of the two parents and one additional fitted parameter; see their Equation 6), matching empirical data sets well. Importantly, Simon *et al.* (2018) take the divergence between the parents as a given (the number of diverged sites and the average trait variance contributed by each site). By contrast, we focus here on how the history of environmental change influences the genetic differences observed between the parents and the amount of hybrid breakdown (i.e., the fitted parameter in Simon *et al.* (2018)). In particular, we explore the conditions under which large effect mutations are likely to accumulate, leading to greater trait variance per locus and greater hybrid breakdown, even among parental populations experiencing the same environments. We then use the Brownian bridge approximation of Simon *et al.* (2018) to fit fitness data from a variety of hybrid types.

Specifically, using simulations, we assess how the history of adaptation to environmental change and the degree of environmental similarity between two populations impact the mean fitness of a variety of hybrid crosses (e.g., F_1_, F_2_, backcross to local, backcross to foreign parents) for both haploid and diploid species. We illustrate how the degree of reproductive isolation depends on the history of adaptation, the mean mutational size, and the cross considered. Our results emphasize how the nature of past selection – both the direction and the speed of selection in each population – has a substantial effect on the amount of hybrid breakdown predicted.

We also consider how the modularity of the genotype-phenotype relationship impacts the development of reproductive isolation under mutation-order speciation. Modularity refers to the observation that genes cluster according to which traits are affected by genetic perturbations. Such clustering is common, for example in the genotype-phenotype maps measured using large gene deletion collections in yeast, nematodes, and mice (Wang *et al.* 2010). Modularity may arise when genes in the same module are subject to similar regulatory networks or have phenotypic effects that are spatially or temporally confined within a particular tissue or cellular component. To vary modularity, we investigate a generalized form of Fisher’s geometric model (FGM), in which the level of pleiotropy can be tuned. The original FGM assumes that a mutation affects all traits under consideration (i.e., universal pleiotropy; fitness landscape is a single hypersphere). Here, we also consider a “multi-sphere model,” where each mutation affects only one module or cluster of traits (modular pleiotropy, Welch and Waxman 2003; Chevin *et al.* 2010; Wagner and Zhang 2011; Hill and Zhang 2012; Martin 2014). Given that we simulate a changing environment affecting the optimum in only one sphere, we predicted that there would be a higher fraction of compatible genetic changes in organisms with greater modularity, because the pleiotropic effects of adaptive mutations and any compensatory mutations would be restricted to the same subset of traits. Thus, we expected a higher degree of parallel evolution, leading to less reproductive isolation, in cases where the environmental change affected the same module. Unexpectedly we found a non-monotonic relationship between the extent of reproductive isolation and the degree of modularity, which we were able to ascribe to a counteracting tendency for mutations to arise more often in different modules, especially late in the divergence process when there are many modules.

Finally, we explore the potential for hybrids to colonize new sites beyond the range of the parental species. Crosses between individuals from two adapted populations can produce a wide variety of hybrid genotypes, potentially allowing hybrids to survive beyond the environmental conditions in which parental genotypes can survive (Albertson and Kocher 2005). We thus explored the range of environments in which hybrids were more likely to establish than parental individuals, potentially contributing to hybrid speciation (i.e., without a change in ploidy: “homoploid hybrid speciation”; Rieseberg 1997).

Overall, our work explores how the nature of environmental change and the modularity of the genetic architecture influence the development of reproductive isolation and the potential for hybrid speciation.

## Model

We first describe the fitness landscape in our model and then describe how we simulate the evolution of reproductive isolation.

### Fitness landscape

Fisher’s model assumes multivariate stabilizing selection around a single optimum. We extend this model by defining an individual’s phenotype as *m* sets of *n*-dimensional vectors, where *m* is the number of spheres and *n* is the number of traits in each sphere. Specifically, the phenotype of an individual in the *i*th sphere is **z**_*i*_ = {*z*_*i*, 1_, *z*_*i*, 2_, …, *z*_*i, n*_}, whose components *z*_*i, j*_ are the phenotypic values for trait *j*; we denote the individual’s phenotype across all spheres as ***Z*** = {***z***_**1**_, ***z***_**2**_, **…**, ***z***_***m***_**}**.

The fitness component based on the set of traits described by the *i*th sphere depends on the distance of the phenotype from the optimum ***o***_*i*_ = {*o*_*i*, 1_, *o*_*i*, 2_, …, *o*_*i, n*_}, measured as the Euclidean distance 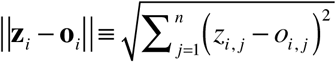. Assuming that an individual’s relative fitness is calculated by considering fitness across all spheres in a multiplicative fashion, fitness is given by:

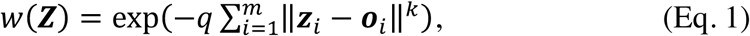

where *q* sets the overall strength of selection and *k* determines the curvature of the fitness function. The parameter *k* corresponds to the degree of epistasis (Tenaillon 2014). Throughout, we assume that fitness declines from a peak height at the optimum, according to a Gaussian fitness surface (*k* = 2). It is worth noting that selection is assumed independent and equally strong on all *m* × *n* traits, for a given distance to the optimum (isotropic selection).

Each mutation additively shifts the phenotypic value relative to its present value. While each mutation has an additive effect on the phenotype, the effect on fitness is not additive but depends on the height of the fitness function for those phenotypes (Eq. 1). In the multi-sphere model, we assume that a mutation affects one, and only one, sphere, chosen at random. Once a sphere is chosen, the mutation affects all of the *n* traits in the sphere under consideration, in a random direction (isotropic mutation). While the trait axes can be defined to allow isotropy with respect to either selection or mutation (assuming a single optimum), it is not possible to do so for both. Nevertheless, previous work exploring a non-isotropic FGM model has shown that the isotropic FGM serves as a good approximation, in many cases, if we consider *n* to be an effective number of dimensions (Waxman and Welch 2005; Martin and Lenormand 2006). Martin (2014) also argued for an isotropic model with universal pleiotropy as an approximation to a broader class of functions.

For both haploids and homozygous diploids, we denote the effect of a mutation occurring within sphere *i* by Δ**z**_*i*_ = {Δ*z*_*i*, 1_, Δ*z*_*i*, 2_, …, Δ*z*_*i, n*_}, with Δ*z*_*i,j*_ describing the effect on trait *j* within that sphere. We assume that a mutation affects each trait within the sphere independently, with Δ*z*_*i, j*_ proportional to a draw from a standard normal distribution, denoted as *ξ*_*j*_ (Hartl and Taubes 1998; Fraïsse *et al.* 2016). The total effect size of the mutation was then set to *r*, drawn from an exponential distribution with mean *λ*, adjusting the mutational effect on each component trait *j* according to 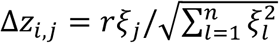 (for alternative distributions, see discussion in Poon & Otto 2000). The phenotypic value of the mutant individual was then set to *z*_*i,j*_ + Δ*z*_*i,j*_ for all traits *j* within sphere *i* and left unchanged for traits in other modules. Denoting the resulting mutant phenotypic vector as ***Z*** + Δ***Z***, the selection coefficient of the mutation is 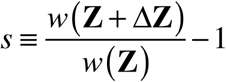. In diploids, the above describes the properties of homozygous mutations. Heterozygous diploids are assumed to be shifted half as far in phenotype space, Δ**Z**/2 (i.e., we assume additivity on a phenotypic scale). The dominance coefficient for fitness is then defined as 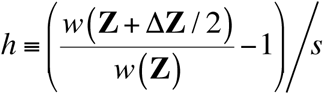.

### Simulation methods

We consider a species that is sexually reproducing with non-overlapping generations. The species is sub-divided into two finite populations, each comprising *N* hermaphrodites with no migration between them (i.e., allopatry). Initially, these populations are genetically identical, having just separated geographically. Genomes may be haploid or diploid and are assumed to be large enough for each mutation to occur at a unique site, with free recombination among sites (i.e., assuming an infinite sites model).

Mutations that affect fitness occur stochastically every generation, but we assume that the mutation rate is quite low to keep the product of the rate and population size much smaller than one. This condition allows us to assume that a population is monomorphic most of the time (Kimura and Ohta 1969; Taylor *et al.* 2006). Thus, we consider the fate of mutations one at a time (Orr 1998; Chevin *et al.* 2014). Each mutation that arises reaches fixation within a single population with probability 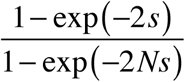 in haploids and 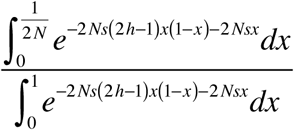 in diploids (Crow and Kimura 1970). Note that overdominance naturally arises, at least transiently, in diploid models of adaptation to a new optimum (Sellis *et al.* 2011). We assume that the population size is small enough that polymorphisms are sufficiently transient to be ignored, although this assumption is more tenuous in diploids with overdominance. To simplify the simulations, we track time in units of fixation events, where we randomly choose mutations and assign them to one or the other population.

As the two populations accumulate mutations, they independently traverse the fitness landscapes in the two patches. While this necessarily leads to genetic divergence, the populations may remain similar phenotypically or may diverge initially and subsequently converge (or vice versa), depending on the optima in the two patches. To explore the impact of parallel versus divergent selection on reproductive isolation, we study the following two environmental scenarios:

1. Adaptation to a common optimum, shifted from the position of the initial population, (to investigate mutation-order speciation).
2. Adaptation to different optima (to investigate ecological speciation).

For each of these scenarios, we explore how the extent of reproductive isolation depends on whether the environment shifts rapidly (in one step) or gradually (in multiple steps). Specifically, we define the vector of optima across all spheres as **O** = {**o**_1_, **o**_2_, …, **o**_*m*_}. By assumption, the initial phenotype ***Z***_0_ and the initial optimum, **O**_0_, are the same in the two populations and set to the origin in all *m* spheres. We then allow environmental change by shifting the optimum phenotype, which induces directional selection until the population approaches the new optimum, at which point the population experiences stabilizing selection. Under scenario (1), the optimum starts at **O**_0_ and moves to a final optimum of **O**_*E*_, either in one step (rapid environmental change) or in ***τ*** time steps, with each step occurring after a fixation event (gradual environmental change). Under scenario (2), adaptation proceeds towards **O**_*A*_ in patch A and towards **O**_*B*_ in patch B, again where the optima move either rapidly once or gradually over ***τ*** fixation events.

We measure the level of reproductive isolation between the two populations through crosses, generating F_1_, F_2_, and backcross (BC) hybrids. To generate F_1_ individuals in the haploid case, for example, when there are *x* fixed mutants in patch A and *y* fixed mutations in patch B, we randomly transmit each mutation to offspring with a 50% probability, so that there are, on average, (*x*+*y*)/2 mutations in each hybrid. In the diploid case, all F_1_ hybrids are identical and are heterozygous for all *x* and *y* mutations. The phenotype of a hybrid individual is then calculated as the sum of the mutational vectors, and its fitness is assessed in both parental environments. This is repeated 1000 times for each hybrid type. FGM simulations for adaptation and for producing hybrids were written in C++ and *Mathematica*, respectively (hosted on Github, https://github.com/ryamaguchi0731/FGM-master).

We were also interested in the time course over which reproductive incompatibilities developed. Whereas some models have assumed that all divergent pairs of loci are equally likely to give rise to speciation barriers (Orr and Turelli 2001), we speculated that large-effect mutations that accumulate during periods of rapid adaptation to a changing environment may contribute disproportionately. For each population, we thus accumulate an equal number of fixation events in two phases: the phase during which adaptation occurs (“adaptation phase”) and the phase during which the population moves randomly around the optimum (“stationary phase”), with drift as well as selection affecting the fate of a mutation. For simplicity, we define the initial adaptation phase as the time until the first mutation fixes with a negative selection coefficient in each population. With a moving optimum, the adaptation phase is defined as the time until the first mutation fixed with a negative selection coefficient after the optimum stops moving, as sometimes deleterious alleles fixed when the phenotype happens to be near the current optimum. In addition, for modular pleiotropy, we assume that the adaptation phase continues until the first allele with a negative selection coefficient fixes within the focal sphere in which the optimum has moved.

To translate the number of fixation events into a more biologically meaningful time scale, we calculated the expected number of generations until appearance and fixation of the next mutation and summed this quantity across the series of mutations fixed. Given the mutation rate *u*, we first calculated the waiting time until the appearance of the next “successful” mutation that would survive stochastic loss while rare. This waiting time was assumed to be exponentially distributed with a rate of events equal to the mutation rate, *u* (set as 10^−5^ throughout this paper) multiplied by the number of haplotypes in the population (*c N*, where *c* = 1 for haploids and 2 for diploids) times the average fixation probability, *P*. The latter varies depending on the current distance of the population from the optimum and was determined by integrating *P* over the distribution of possible mutations for each state of the population. The mean waiting time of this exponential distribution is 1/(*cNuP*) generations. Then, we calculated the fixation time of the successful allele that was drawn, using the expected waiting time until fixation for a new mutation (Ewens 2004; Otto and Day 2007). Given the selection coefficient and the dominance coefficient of a mutation, the expected waiting time, conditional upon fixation, is approximately (Kimura 1980):

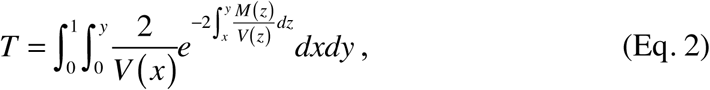

where *M* and *V* are the mean and variance of the change of mutation allele frequency *p* per generation. Specifically, *M* (*p*) = *sp*(1 − *p*) and 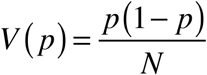 in haploids and *M* (*p*) = *p*(1 − *p*)[*sp* + *h*(1 − 2 *p*] and 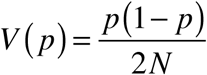 in diploids. The expected waiting time until the next mutation appears and fixes then becomes 1/(*cNuP*) plus *T*.

## Results

### Evolutionary trajectory of adaptation

Under Fisher’s geometric model (FGM), mutations have nearly a 50% chance of being beneficial when a population starts far from the optimum relative to the effect size of the mutation, but they are more likely to overshoot the optimum and less likely to fix once the population approaches the optimum (Fisher 1930; Kimura 1983). As a consequence, during the process of adaptation, the average phenotypic effect size of fixed mutations is typically larger at first and declines as populations become increasingly adapted (Figure S1a; Orr 1998). When the distribution of mutations has a larger average magnitude, we find that the optimum is reached faster, as expected (Figure S1c). After nearing the optimum, populations accumulate a combination of slightly deleterious and beneficial mutations (inset to Figure S1b), resulting in a phase of mutation-selection-drift where the populations stochastically move around the optimum through successive fixation events.

We then accumulated the same total number of fixation events in the initial adaptation phase and in the second stationary phase in each parental population (allowing each parental population to accumulate a different number of fixed mutations). This procedure ensured that we allowed enough time to pass in the stationary phase that the level of divergence contributed by the two phases is the same (in terms of substitutions). Table S2 gives the number of fixation events in each parental population, the corresponding expected number of generations, and the proportion of time in the adaptation phase. The adaptation phase was typically much faster than the stationary phase. After the adaptation and stationary phase had finished, we performed crosses between populations to determine how hybrid fitness depended on the nature of environmental change, the mutation effect size, ploidy, and the modularity of pleiotropy.

### Adaptation to a common optimum

Figure 2 shows hybrid fitness after a period of adaptation towards a common optimum in haploids and diploids (for an exploration of introgression effects, see Fraïsse *et al.*, 2016), assuming that the optimum shifted in one step (rapid environmental change). For the parameters considered, substantial intrinsic reproductive isolation is observed in all hybrids, with the exception of diploid F_1_s. Diploid F_1_s are not yet recombinant but heterozygous at all sites that differ between the parents. When parents near the same optimum are crossed in the FGM, averaging their phenotype in the diploid F_1_ reduces the distance to the peak, as discussed by Barton (2001), Fraïsse et al. (2016), and Simon et al. (2018). Indeed, we find that the fitness of F_1_ hybrids is about three times closer to the optimum than the average parental fitness, although this difference is too small to see in Figure 2 (parental and F_1_ hybrid fitnesses of 0.999273 and 0.999784, respectively, in panel (b) with *λ*= 0.04 and of 0.997224 and 0.999152 in panel (d) with *λ*= 0.12).

By contrast, all recombinant hybrids have reduced average fitness, relative to the parents, reflecting the segregation load studied by Chevin *et al.* (2014). Importantly, the incompatibility results primarily from the large-effect mutations that accumulate during adaptation, which combined can generate hybrids that move far off the peak fitness and that exhibit substantial pleiotropic side effects (Figure 1). Such incompatibilities are not observed to any substantial degree when the optimum remains at the origin, even with the same number of fixation events (Figure S2). That is, reproductive isolation is not simply a function of the number of divergent alleles. Rather, the vast majority of reproductive isolation arose during the first adaptation phase when large-effect mutations were likely to fix. Considering only these mutations, F_2_ hybrid fitness, for example, was reduced below the optimum by, on average, 0.72 in haploids and 0.74 in diploids. By contrast, F_2_ hybrid fitness remained much higher, 0.99 in haploids and 0.98 in diploids, when we add the same number of mutations that occurred only during the stationary phase to the optimum (see Table S2 for the number of fixation events in each population).

Hybrid breakdown did not accumulate linearly over time during the adaptation phase (Figure S3a). Indeed, early on hybrids were slightly fitter than the average of the parental populations when fitness was measured in their new environment, presumably due to the large advantage of combining mutations and approaching the distant optimum. Only after several mutations had fixed did hybrid fitness decline, on average, relative to the parents.

We next asked how much of the hybrid breakdown was caused by mutational effects along the trait axis of environmental change (e.g., hybrids overshooting the optimum) versus pleiotropy (e.g., hybrids breaking apart compensatory gene combinations). To do this, we measured hybrid fitness only along the axis of environmental change, setting all other trait values to the optimum (black slice in pie chart). The remainder of the hybrid breakdown is due to pleiotropy. We find that the majority of the observed breakdown was due to pleiotropy (Figure 2).

While we have illustrated the fitness drop in several hybrid forms, the Brownian bridge approximation of Simon et al. (2018) allows us to connect the fitness of any hybrid genotype, given the parental fitness, the hybrid index (*H*), the heterozygosity of the hybrid (*p*_12_), and one free parameter, which measures the extent of hybrid breakdown. As illustrated in Figure S4 (a,b), the method of Simon et al. (2018) works well to predict the fitness of each class of hybrids, once we fit the free parameter by minimizing the sum of squared deviations between the observed and expected fitnesses for the different hybrids.

During the period of adaptation, large mutations can fix, even if they shift the phenotype substantially away from the origin in some directions, as long as the average phenotypic distance to the optimum is shorter. In subsequent generations, these negative pleiotropic side effects favor the fixation of compensatory mutations that move the phenotype back toward the optimum (Figure 1). In recombinant hybrids, however, these compensatory mutations are separated from the mutant backgrounds in which they are favored, reducing fitness. Consequently, the fitness reduction in hybrids is more severe when mutational effect sizes (and hence pleiotropic side effects) are larger, on average (see Figure 3a,b with a 0°angle for populations experiencing a common optimum, with dots from left to right representing smaller to larger average effects). It is also worth highlighting the substantial amount of variation in reproductive isolation observed across replicates, depending on the exact adaptive path taken, with more variability occurring when the average mutational effect is larger (for Figure 3a, for example, the standard deviation in the average F_1_ haploid fitness among replicates rose from 7.41×10^−3^ when *λ*=0.04 to 1.26×10^−2^ when *λ*=0.12).

**Figure 3.**
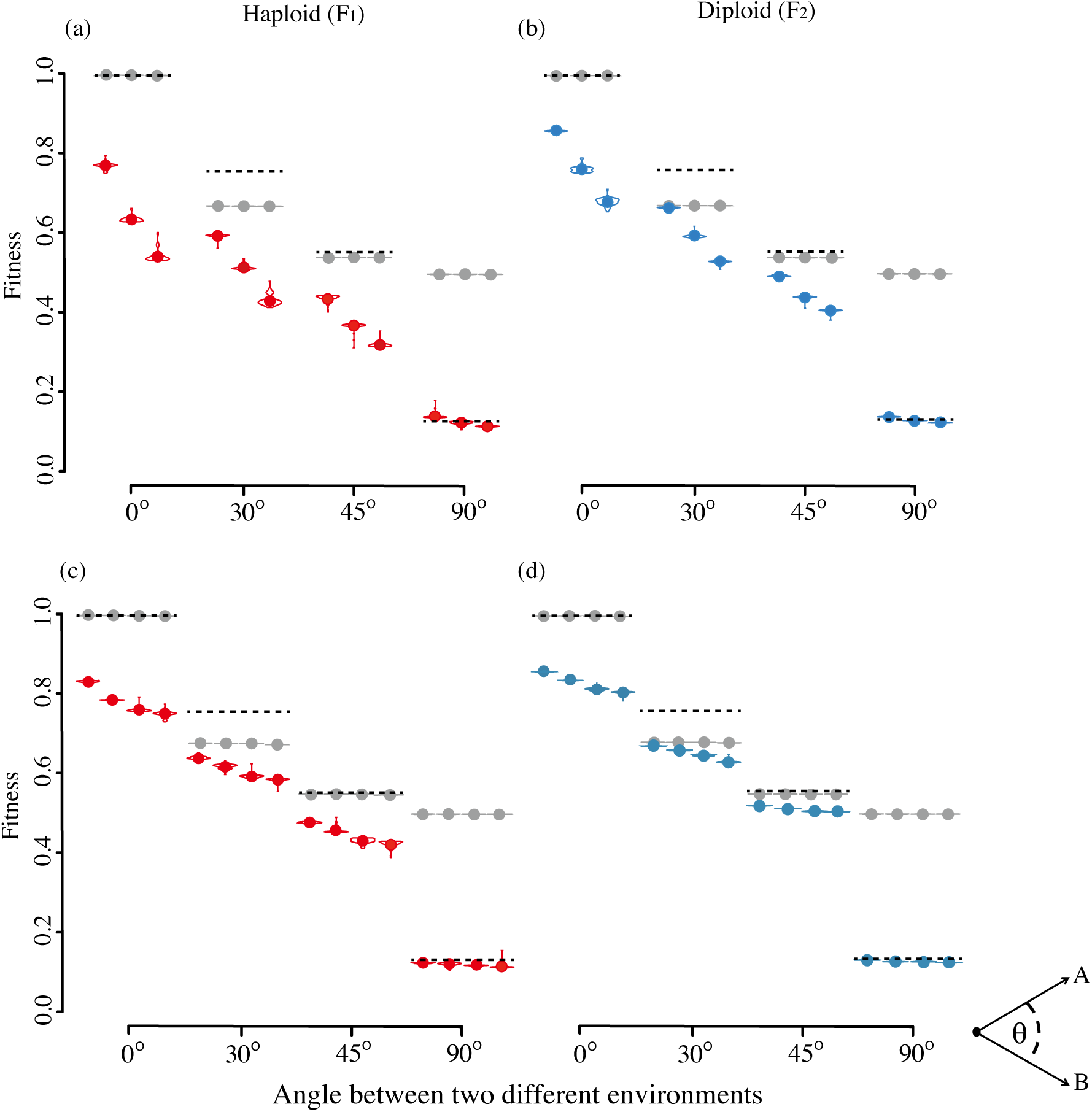
Stronger reproductive isolation arises via ecological speciation when populations are selected in more divergent directions, with more isolation arising when mutations have a larger average effect size (top panels) and the environment changes rapidly (bottom panels) in haploids (a, c) and diploids (b, d). For top panels, mutations are drawn from an exponential distribution with *λ* = 0.04, 0.08, and 0.12 (three adjacent points from left to right). For bottom panels, the optimum is gradually moved from the origin in one trait dimension over *τ* = 100 (slow), 50 (medium), 20 (fast) or 0 (immediate) fixation events until the new optimum is reached (four adjacent points from left to right), assuming *λ* = 0.04. The x-axis gives the angle between the optima **O**_*A*_ in patch A and **O**_*B*_ in patch B relative to the ancestral optimum **O**_0_ (as illustrated on the right bottom). Grey points show the average fitness of the two parental genotypes (we arbitrarily measure all fitnesses in patch A, to which population 1 is adapted). Dashed lines show the expected fitness of the hybrids at the parental mid-point. All other parameters are as in Figure 2 except that this figure is calculated from 50 simulations.

The above simulations were repeated but allowing the optimum to move gradually towards its final position, **O**_*E*_. With a history of slower environmental change, large-effect mutations have a lower probability of fixation, reducing the accumulation of negative pleiotropic side-effects. Consequently, we find that much weaker reproductive isolation develops, with higher hybrid fitness, after a period of slow environmental change, even when ultimately reaching the same optimum (see Figure 3c,d with a 0°angle for populations experiencing a common optimum, with dots from left to right representing slower to faster environmental change). We thus conclude that rapid environmental change, resulting in the accumulation of large-effect mutations, is particularly conducive to mutation-order speciation.

### Adaptation to different optima

In the next scenario, we simulated isolated populations adapting to different fixed optima, assessing hybrid fitness in both parental environments. The results are qualitatively different in diploids, and now all hybrids, including F_1_ hybrids, have lower fitness within each environment, as their phenotype falls between the optima of the two environments (extrinsic reproductive incompatibility). The larger the angle between the two optima, the lower the fitness of hybrids (Figure 3), as well as the lower the average fitness of the two parents (grey dots). The average fitness of the two parents is very close to that expected by the respective optima to which they have adapted (Table S1), with one parent having fitness near one and the other having a fitness that declines to zero as the angle of divergence rises. Also shown is the expected fitness if the hybrid were phenotypically mid-way between the two parents (dashed line). When the new environmental optima are at 30°or 45°relative to the ancestral population, the recombinant hybrids are less fit than either the average of the parents or the parental mid-point, indicating intrinsic incompatibilities. At 90°, some of the recombinant hybrids have a high proportion of local alleles, causing the hybrid offspring to rise above that expected for the mid-parent phenotype (less extrinsic incompatibility). Hybrid breakdown is more severe when mutational effect sizes are larger (see sets of three adjacent points in Figure 3a,b), indicating stronger intrinsic incompatibilities, but this effect is weak when parents are highly diverged (90°) because of the counteracting advantage of recombinants being more likely to carry large effect alleles adapted to patch A. The temporal dynamics of hybrid breakdown is illustrated in Figure S3. Again, divergence accumulates slowly at first, especially when the new optima are similar (30°or 45°).

We next explored the sensitivity of ecological speciation to the speed of environmental change. As observed with mutation-order speciation, greater reproductive isolation arises when the environment changes rapidly (see sets of four adjacent points in Figure 3c,d), but again the effect is smaller than the effect of the angle between the two environments. Essentially, the strong extrinsic reproductive incompatibility caused when parents are adapted to different environmental optima masks the intrinsic incompatibilities that arise from accumulating incompatible mutations (Figure 1). The more similar the current parental environments (the lower the angle between them), the greater the difference in hybrid breakdown observed between a history of slow versus rapid environmental change.

Again, the Brownian bridge approximation of Simon et al. (2018) allows us to correctly predict the fitness of any hybrid genotype, by fitting one parameter along with the parental fitnesses, *H*, and *p*_12_ (Figure S4). The fitted parameter, a measure of the extent of hybrid breakdown, depends on the current parental environments and is greater when populations have been selected to more divergent optima (compare Figure S4 panels c,d to e,f). Consistent with the conclusions drawn above, the breakdown parameter is also greater when fit to simulation data that have allowed larger-effect mutations to accumulate, either because the average effect of mutations is larger (as in Figure 3a,b) or because the environmental change occurred faster (as in Figure 3c,d).

When parental population have experienced divergent selection toward different optima, backcross hybrids to the local parent, BC_L_, show much higher fitness than backcross hybrids to the foreign parent, BC_F_ (Figure S5a,b), a hallmark of ecological speciation (Rundle and Whitlock 2001). The relative performance of the various hybrids depends, however, on the angle between the two optima (compare Figure S5c,d with an angle of 90°to Figure S5a,b with 30°). As a measure of the importance of ecological speciation, Rundle and Whitlock (2001) suggest calculating the fitness of BC_L_ minus the fitness of BC_F_, averaged over the two local environments, as this estimates the strength of additive-by-environment interactions (AE). Indeed, AE rises from zero when parental environments are identical to ∼0.4-0.6 for environments separated by an angle of 90°(Figure 4a,b). Across the full range of angles, however, AE behaves non-monotonically (Figure 4a,b). For very dissimilar environments, the fitness of the local BC_L_ falls to zero, severely limiting the maximum value of AE.

**Figure 4.**
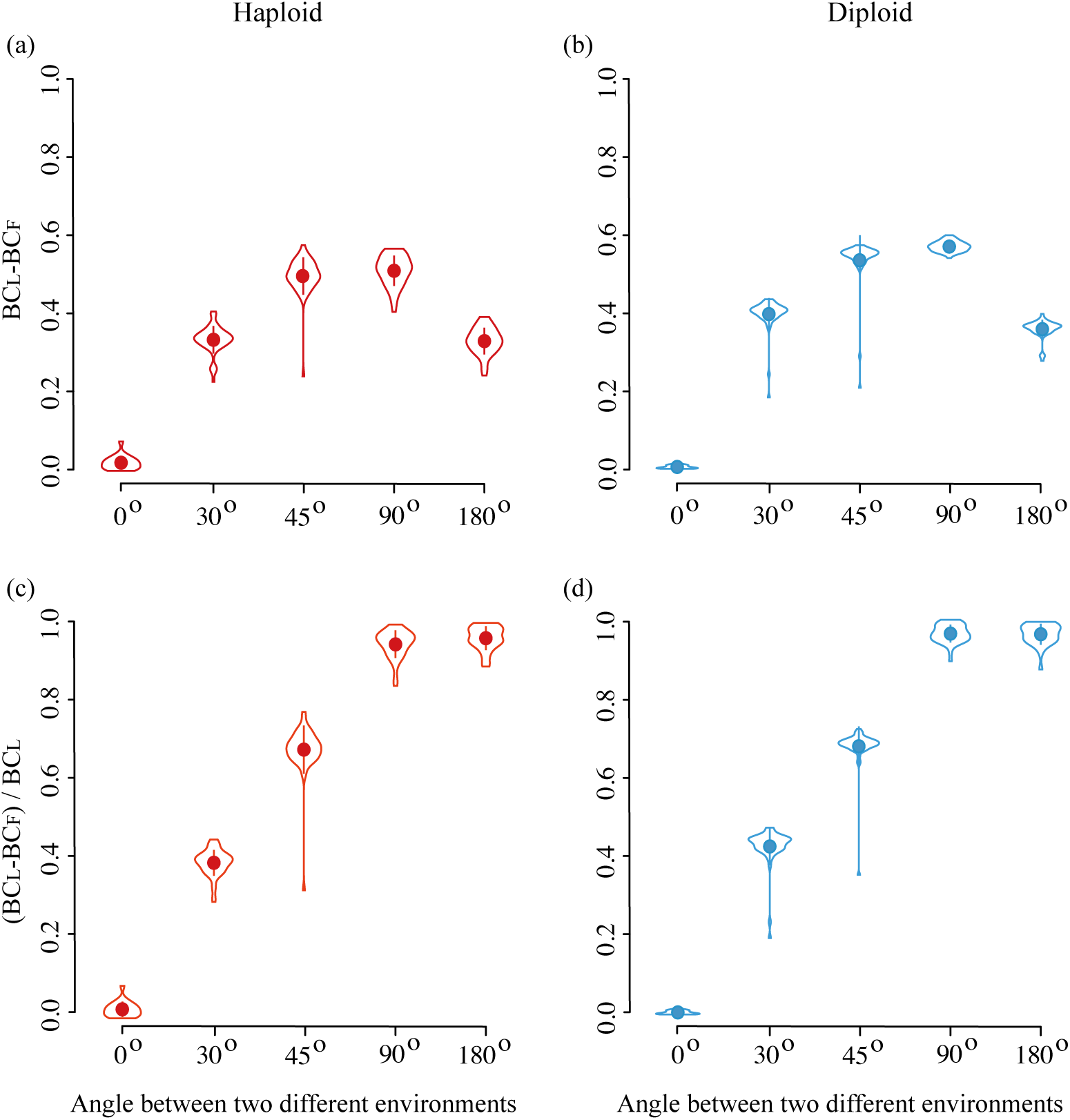
Estimating the degree of ecologically-dependent reproductive isolation. Top panels give BC_L_ – BC_F_ for (a) haploids and (b) diploids, as a function of the angle between the optima toward which the parental populations were evolving. Bottom panels give (BC_L_ – BC_F_)/BC_L_ for (c) haploids and (d) diploids. While BC_L_ – BC_F_ shows a non-monotonic relationship, (BC_L_ – BC_F_)/BC_L_ rises monotonically with the degree of ecological difference between the two environments. Mutations are drawn from an exponential distribution with *λ* = 0.04. All other parameters are as in Figure 2, with distributions and solid points representing the mean ± SE calculated from 50 simulations.

The degree to which ecological speciation contributes to reproductive isolation may be better captured by measuring AE relative to the maximum value it could take (i.e., AE divided by the fitness of BC_L_). This rescaled value measures the degree to which additive-by-environment interactions reduce the fitness of hybrids backcrossed to the foreign parent compared to the fitness expected based on backcrosses with the local parent. When AE/BC_L_ is zero, fitness is the same regardless of which parental population is used in the backcrosses (as expected when the parental environments are the same), whereas an AE/BC_L_ of one indicates that the genetic differences between the parents reduces the fitness of backcross hybrids to zero when raised in the foreign environment (as expected following strong divergent selection). The rescaled measure AE/BC_L_ does rise monotonically with the angle between the optima and hence the degree of divergent selection (Figure 4c,d), although sample sizes would need to be sufficient to avoid bias caused by measurement error in the denominator.

### Impact of modularity

We next investigated the degree to which modularity of the phenotype would affect the development of reproductive incompatibilities between adapting populations. To investigate different degrees of modular pleiotropy, we held the total number of phenotypic dimensions constant (the number of spheres times the number of traits per sphere = *m n*) but varied the number of spheres. To simplify the presentation, we consider only F_1_ hybrids in haploid simulations. Our prediction was that we would observe less negative pleiotropy when the phenotype was more modular, leading to less hybrid breakdown. While we did see that, as expected, pleiotropy contributed less to hybrid breakdown in more modular simulations (pie charts in Figure 5a), hybrid fitness actually declined with high levels of modularity (Figure 5a). Specifically, reproductive isolation showed a non-monotonic pattern, with more hybrid breakdown in the fully pleiotropic model (*m* = 1) and in the model with no pleiotropy (*n* = 1) than with intermediate levels of pleiotropy.

**Figure 5.**
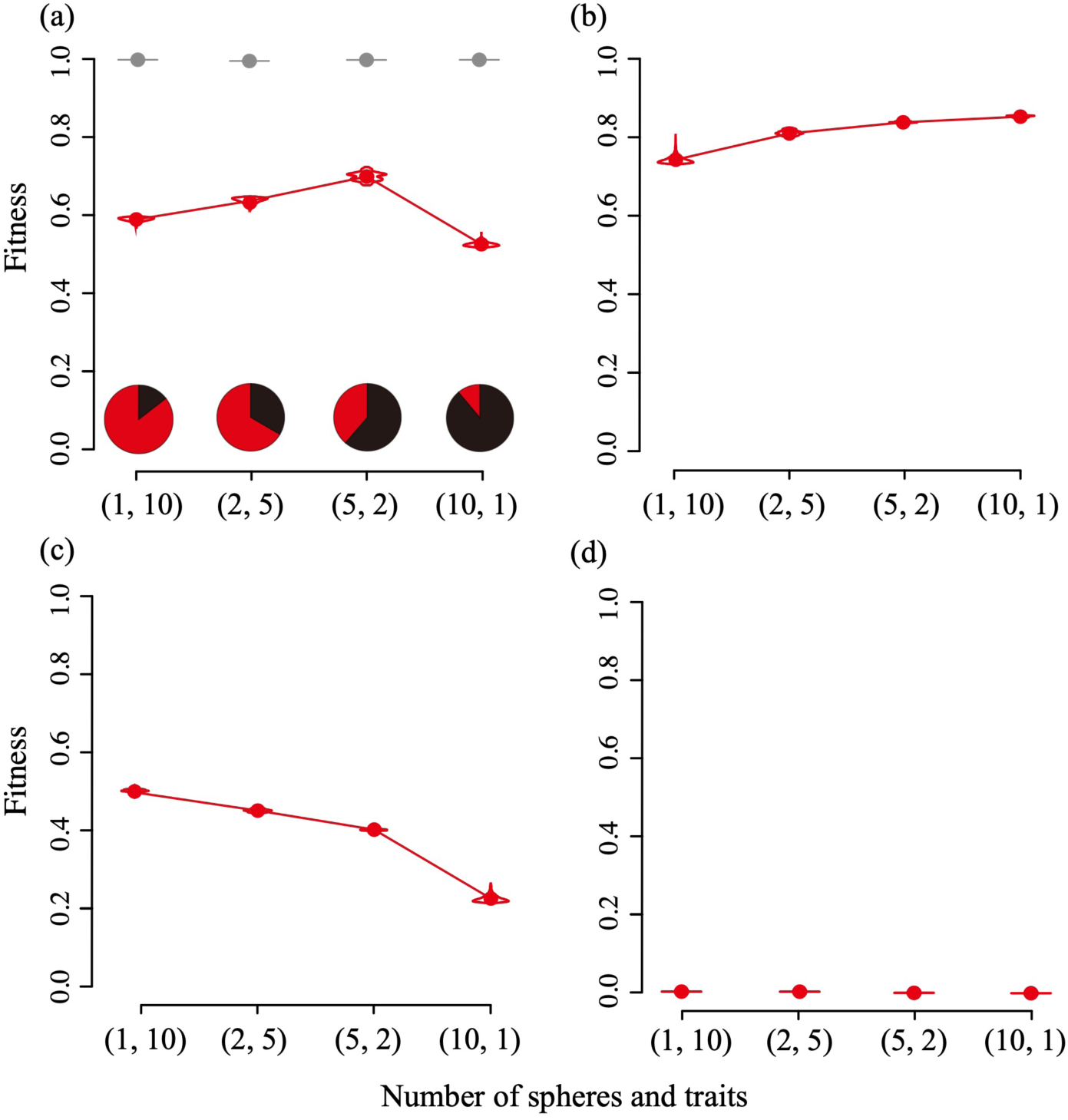
Effect of modularity on the accumulation of reproductive isolation in haploid F_1_ hybrids evolving to a common optimum. Panel (a) includes all mutations fixed, while (b) includes only the first 50% of mutations fixed during the adaptation phase and (c) includes only the second 50% of mutations fixed during the adaptation phase; panel (d) includes only mutations fixed during the stationary phase. In each case, the phenotypic effects were added to parents whose phenotypes were at the origin. In (a), pie charts illustrate the fraction of the total hybrid breakdown along the single phenotypic axis experiencing environmental change (black), versus fitness effects on all other trait dimensions. All other parameters are as in Figure 2 with *λ* = 0.04, except that this figure changes the number of spheres along the x-axis (from 1 on left to 10 on right) and is calculated from 50 simulations.

To understand this counter-intuitive result, we investigated how the genes contributing to hybrid breakdown accumulated over time. Relatively little contribution came from mutations accumulating during the stationary phase, so we sub-divided the mutations accumulated during the adaptation phase into two groups: the first 50% of mutations that fixed and the second 50% of mutations that fixed. With more modular genetic networks (*m* larger; Figure 5b), mutations that fixed during the first half of adaptation were more likely to work well together, improving fitness along the axis of environmental change and increasing mean hybrid fitness, as we had expected. When the environmental change occurs in a module with few traits (low *n*), mutations that fix in one population are more likely to point towards the optimum and still be beneficial in another population. By contrast, as the population approached the optimum during the second half of the adaptation phase, small-effect mutations in different modules made up an increasingly large fraction of the mutations that fixed in different populations (recall that the stationary phase was defined to start only after a deleterious mutation arose in the module experiencing environmental change). Consequently, the more modules, the less likely mutations in different populations would compensate for one another, so that decreased hybrid fitness was observed with more modularity (Figure 5c). Thus, the role of modularity in speciation had opposite effects during rapid early periods of adaptation, where reproductive isolation developed faster in less modular systems, compared to later periods of relatively slow adaptation, where reproductive isolation developed faster in more modular systems.

As an alternative explanation, we asked whether there was a substantial difference in the magnitude of mutations as a function of modularity. The magnitude of the largest mutation that fixed in each population was, however, similar across the combinations of *m* and *n* explored (Figure S6), and so this explanation does not appear to contribute substantially to the non-monotonic pattern with modularity.

### Homoploid hybrid speciation

As discussed above, the history of environmental change can have a strong impact on the degree of reproductive isolation exhibited among populations. Here, we briefly explore how this history can also impact the potential to produce hybrids that are able to colonize new environments, leading to hybrid speciation. Specifically, we investigated the possibility of homoploid hybrid speciation by crossing the two parental populations generated in our simulations and measuring the fitnesses of hybrids across a range of possible new environments. For each environment, representing a potential site for colonization, the fitness of F_1_ hybrid haploids was assessed given the optimal phenotype in that environment, **O**_*P*_. The probability of establishment in a site, *P*_e_, was determined for any genotype by the branching process approximation 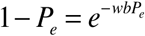, assuming a Poisson number of offspring with mean equal to the genotypic fitness *w* times the average reproductive capacity, *b*, of an individual with full fitness of *w*=1 (*b* was set to 5 in the simulations). We repeated this calculation for 1000 hybrids and averaged *P*_*e*_ to get the expected establishment probability. We then measured the establishment probability of hybrids relative to the average establishment probability of the two parental genotypes.

Figure 6 illustrates the colonization success of hybrid haploids versus their parents, as a function of the phenotypic optimum in the colonized environment. Whether hybrids are more likely to colonize a given environment depends strongly on the history of adaptation in the parental populations. When the parental populations diverged in a constant environment, hybrids are almost never better colonists than the parental populations (top panels; optimum constant at zero as in Figure S2). By contrast, hybrids are more variable following a period of adaptation to a common environment (mutation-order speciation), and so at least some individual hybrids are able to colonize substantially different environments than the parental populations (using parents from simulations reported in Figure 3 with a 0°angle between environments). With ecological speciation between parental populations adapted to different environments (> 0°angle in Figure 3), hybrids are much better at colonizing new environments that fall between the optima of the parental populations (bottom panels). Homozygous diploid F_2_s behave similarly, but the establishment probability of diploids in general would have to account for segregation variance after colonization, which was not explored.

**Figure 6.**
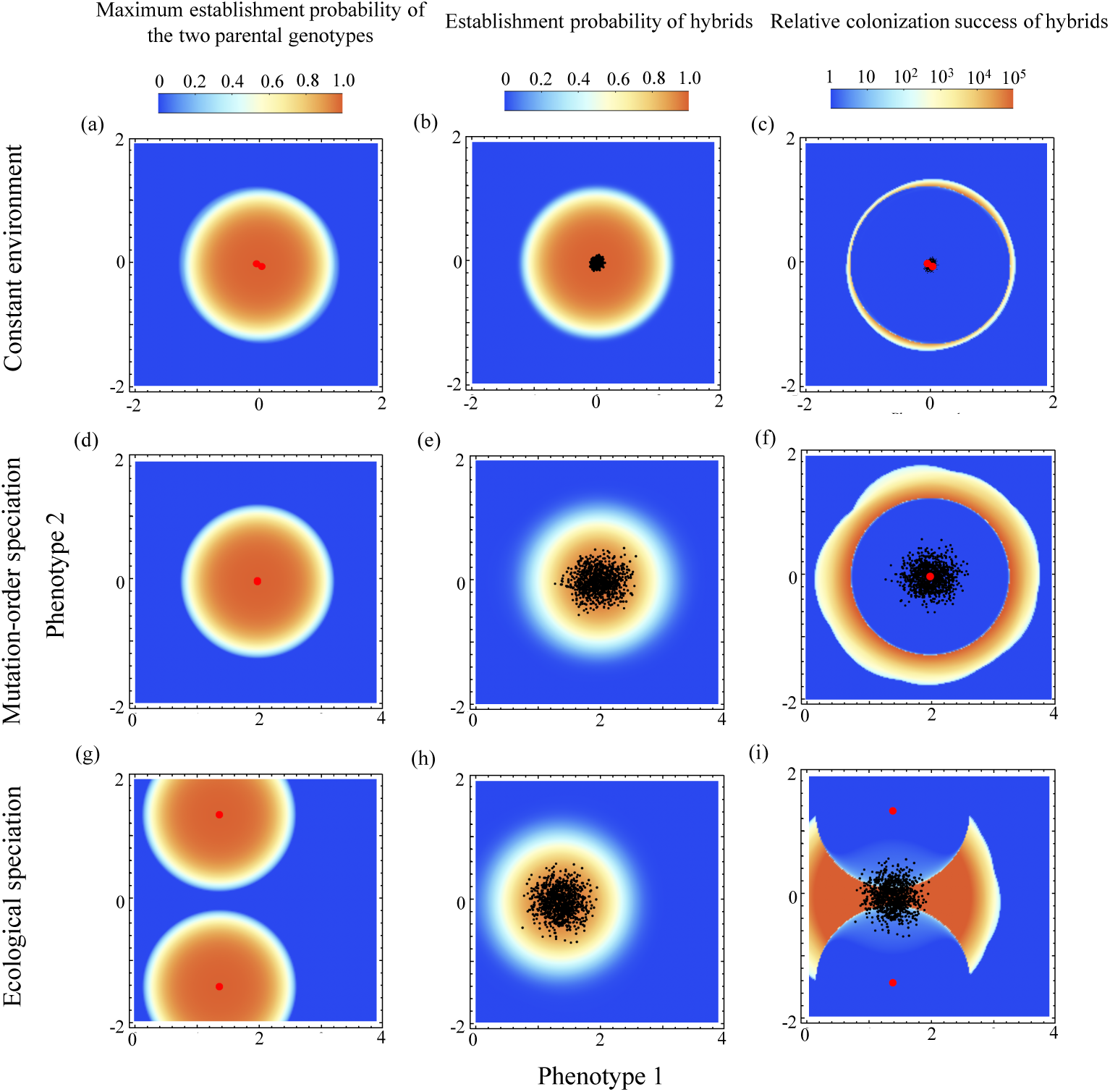
Establishment probability of haploid hybrids relative to parents when colonizing a new environment. The optimal phenotype in the new environment is varied along two dimensions (x and y axes), holding the optimum at zero for all other phenotypic dimensions (*n*=10). Left panels show the establishment probability measured in each environment for the best of the two parental genotypes (red dots). Middle panels show the establishment probability of F_1_ recombinant hybrids (black dots). Shading indicates establishment probability, from low (blue) to high (orange), as indicated at top. Right panels show the colonization success of hybrids relative to the parents, where the degree of orange indicates the amount by which hybrids have a mean establishment probability greater than the mean of the two parents (fixation probabilities <10^−6^ were set to 10^−6^ to avoid division by small numbers). The potential for hybrid speciation depends strongly on the history of the parental populations: (a-c) diverged in a constant environment (parents from Figure S2a), (d-f) diverged through parallel adaptation to a new optimum at 2 along the x-axis (mutation-order speciation, parents from Figure 2a), (g-i) diverged through adaptation to different environments (ecological speciation with an angle between the two optima of 90°, parents from Figure S5c). All other parameters are as in Figure 2, with *b* = 5 and *λ* = 0.04.

Thus, the history of the parental populations has a large influence on the fate of hybrids and their ability to survive in and colonize new environments. Even when the parental populations are phenotypically similar, distinct hybrids that can survive quite different environmental conditions are more likely to be produced if the parents had recently undergone a period of adaptation and accumulated different large effect mutations (compare middle panels to top panels of Figure 6).

## Discussion

We used a simple model of optimizing selection on multiple traits (“Fisher’s geometric model”) to explore speciation in the face of environmental change. Because the path of adaptation in Fisher’s geometric model varies among populations evolving in allopatry, genetic crosses between populations can yield mis-matched combinations of adaptive mutations, producing hybrid offspring of lower fitness (Chevin *et al.* 2014; Fraïsse *et al.* 2016; Simon *et al.* 2018). Our main conclusion from this model is that periods of environmental change are likely engines of speciation for allopatric populations, allowing populations to develop strong intrinsic reproductive incompatibilities even when they are adapting to similar environmental changes (“mutation-order speciation”; Figure 3), as well as contributing additional intrinsic incompatibilities to the extrinsic hybrid breakdown that occurs when parents are adapted to different environments (“ecological speciation”; Figure 3). Reproductive incompatibilities were strongest when large-effect mutations could accumulate, i.e., during times of rapid environmental change, which challenges the notion that all genetic substitutions are equally likely to generate reproductive incompatibilities.

Using the metaphor of a “speciation clock” (e.g., Coyne & Orr 1989), these results imply that the clock will vary in rate, sometimes speeding up and sometimes slowing, depending on the local environmental circumstances and the chance of experiencing a period of adaptation. Averaged over these circumstances, however, we would still predict a constant rate of ticking, but our work predicts greater variation in the amount of BDMIs for a given period of divergence time, depending on the history of stasis or change in the environment.

We also explored the relative importance of mutation-order speciation versus ecological speciation by comparing the fitness of different hybrid crosses. Rundle and Whitlock (2001) showed that the contribution of ecological speciation could be captured by the additive-by-environment interaction (AE), estimated from the fitness of individuals backcrossed to the local population (BC_L_) minus the fitness of individuals backcrossed to a foreign population (BC_F_). Only under ecological speciation are these backcross fitnesses expected to differ. We find that this AE measure does indeed rise with the angle of environmental optima between two evolving populations (measured here relative to their ancestral origin), but only when that angle was small. With larger environmental differences, the fitness of even the local backcrossed individuals (BC_L_) could be very low, leading AE to drop (Figure 4a,b). We thus caution against using low AE values to distinguish cases of mutation-order from ecological speciation, especially when backcross fitnesses are all low. As a correction for this non-monotonic behavior, we instead investigated a new measure of the importance of ecological speciation, dividing AE by its maximum expected value (BC_L_), which we found does rise with the angle of environmental differences in our study and hence is a better indicator of the extent of divergent selection (Figure 4c,d). It would be valuable for future studies to determine the sampling properties of this new measure and its ability to distinguish mutation-order and ecological speciation in simulated and empirical studies.

In addition, we considered how the modularity of the genotype-phenotype relationship impacted the development of reproductive isolation under mutation-order speciation. With a more modular genetic architecture, allopatric populations experiencing the same environmental change are more likely to accumulate mutations that are compatible, because the negative pleiotropic effects of adaptive mutations are constrained within the module affected by the environmental change. Thus, we expected a higher degree of parallel evolution, leading to less reproductive isolation, in cases where the environment changed in the same direction in different populations. Unexpectedly, we found a non-monotonic relationship between the extent of reproductive isolation and the degree of modularity (Figure 5). By separating out the first half of mutations fixed during the rapid early phase of adaptation from the latter half of mutations fixed during the adaptation phase, we were able to verify that reproductive isolation is less likely to develop in more modular systems early on during adaptation when selection is strong, but this is counteracted by a greater chance that mutations generate reproductive isolation during later phases of adaptation as populations more slowly approach the optimum. The key difference between these phases was whether fixed mutations tended to fix primarily in the same module – the sphere with the shifting environment – or in different modules. Once the populations were near the optimum, different populations tended to fix mutations and compensatory mutations in different modules, substantially increasing the chance of hybrid breakdown when crossed.

Finally, we explored how the history of adaptation of the parental populations influenced the potential to produce hybrid offspring that could colonize a new environment and lead to homoploid hybrid speciation. Because hybrids can vary in phenotype beyond the range of parental populations, hybrids can successfully colonize environments inaccessible to parental populations, potentially contributing to homoploid hybrid speciation (Gross and Rieseberg 2005). Importantly, we found that hybrid speciation was especially likely when the parental populations had undergone a period of rapid adaptation in allopatry (Figure 6). The parents were then more likely to differ by large-effect mutations, allowing hybrid genotypes to vary more broadly in phenotype and producing at least some individuals that could succeed in more extreme environments. The potential for colonization success by hybrids was observed whether the parental populations had adapted to a common environment (mutation-order speciation, Figure 6f) or different environments (ecological speciation, Figure 6i), although the environment where hybrids were most successful differed (in a halo around the common parental environment versus midway between them, respectively). We thus predict that periods of adaptation by the parental forms may be particularly conducive to subsequent homoploid hybrid speciation.

Recombining pre-existing variation is a powerful way of generating new species over short periods of time. Indeed, hybrid speciation is thought to underlie adaptive radiation in a number of systems (Marques *et al.* 2019; Martin and Richards 2019). Based on our simulations, we suggest that large effect mutations following a period of adaptation may also contribute disproportionately to admixture variation, contributing to intrinsic and extrinsic incompatibilities, transgressive traits, and novel trait combinations in hybrid swarms (compare, for example, figure 6f,i to 6c).

In this study, we have made several simplifying assumptions that warrant more exploration in future work. For one, we assumed that populations are largely monomorphic, punctuated in time by the fixation of new mutations. This assumption breaks down in larger populations (*N*) or in environments with a larger number of potential mutations (higher *u*), either of which increases the flux of mutations into a population. A higher flux of mutations can increase the rate at which large-effect mutations appear, increasing the probability that intrinsic incompatibilities arise, but more mutations also increase the standing genetic variation of the parental populations, which can mean that the same alleles are present and fix in the parental populations when the environment changes (Thompson *et al.* 2019). Which effect is more important depends on the exact details, e.g., when the parental populations became isolated relative to the change in environment. Future work is needed to explore the development of reproductive isolation allowing for polymorphic alleles at multiple loci throughout the genome. Relaxing the assumption of monomorphism is particularly important for diploid populations, because large effect mutations often arise that overshoot the optimum, leading to overdominance that maintains polymorphism (Sellis *et al.* 2011).

Another simplifying assumption in our model is the Gaussian fitness landscape (*k* = 2). Fraïsse *et al.* (2016) investigated various *k* values with divergence via genetic drift around an optimum and found that speciation genes could fix with a nonnegligible probability only when the number of traits (*n*) was sufficiently small and *k* was sufficiently large (their Figure 2a). This is consistent with our finding that BDMIs do not develop to any appreciable extent in populations that remain near the optimum. With a history of adaptation allowing large-effect mutations, however, BDMIs do arise even with a quadratic fitness surface (Figure 2 here, see also Barton *et al.* 2001), although the results of Fraïsse *et al.* (2016) suggest that even more hybrid breakdown would be observed with *k* > 2.

In addition, it would be worthwhile exploring other forms of selection beyond the single-peaked surface considered here. With multiple optima or other complex fitness landscapes (e.g., the “holey adaptive landscape” of Gavrilets 1997), mutation-order speciation is thought to be even more likely, as populations move to different points on the fitness landscape, depending on the order in which mutations arise (Gavrilets 2004). Nevertheless, we conjecture that the degree of reproductive isolation may still be stronger when initially further away from the peaks of a rugged fitness surface because large-effect mutations may then be more likely to fix in allopatric populations, relative to the small-effect mutations providing slight improvements in fitness near a local optimum. Future work would be valuable that explores how the shape of the fitness landscape affects the likelihood of accumulating the large effect mutations that contribute disproportionately to hybrid breakdown.

As an example of speciation across a more complex fitness landscape, Benkman (2003) showed that the fitness surface of red crossbills feeding on cones from different coniferous tree species displayed different peaks, associated with distinct call types, with both ridges of relatively high fitness and deep fitness valleys. Although not explored in this paper, the combination of fitness ridges with deep valleys might facilitate reproductive isolation by allowing large-effect mutations that reach different points along the ridge and that, when combined, cause hybrid phenotypes to exhibit low fitness.

Fluctuating selection could also contribute to the accumulation of large-effect mutations and be a potent driver of speciation (see, e.g, Barton 2001; Chevin *et al.* 2014; Fraïsse *et al.* 2016). For example, beak phenotypes of the medium ground finch *Geospiza fortis* on Galapagos have experienced selection that fluctuates between life stages, years and generations (Price and Grant 1984; Grant and Grant 2008). Stabilizing selection with a fluctuating optimum allows the population to accumulate relatively large mutations, which contribute disproportionately to the accumulation of reproductive isolation.

In conclusion, our work shows that the tempo of speciation does not tick at a steady rate. Rather, reproductive isolation between allopatric populations should rise faster after bouts of rapid environmental change. A rapidly changing environment is particularly important when selection acts in parallel among populations (mutation-order speciation), as extrinsic reproductive incompatibilities are lacking. In our simulations, crossing the few large-effect mutations that arose during bouts of rapid adaptation caused more hybrid breakdown than combining many small-effect mutations, whose effects tended to cancel (Figure 1; see also eq. 3 in Chevin *et al.* 2014). These results help explain the surprisingly rapid appearance of genetic incompatibilities during short-term experimental evolution studies in the lab (e.g., Dettman *et al.* 2007, 2008; Kvitek and Sherlock 2011; Chou *et al.* 2014; Ono *et al.* 2017). Periods of strong selection are unusually powerful drivers of speciation, helping to account for why reproductive isolation arises so rapidly in experimental evolution studies and yet the biological world has not diversified into an infinite variety of species.

## Author contributions

RY and SPO designed the study; RY developed the computer simulation code and conducted analyses in consultation with SPO; RY and SPO wrote the manuscript.

## Data accessibility

All code necessary to repeat the analysis described in this study have been made available. C++ source codes of our simulation models for adaptation process, *Mathematica* (version 11.2.0.0) codes for calculating hybrids genotypes and fitnesses, R (version 3.5.1) scripts that we used to produce figures, and simulation results in this paper are hosted on Github (https://github.com/ryamaguchi0731/FGM-master).

## Acknowledgements

This work was supported by funding from the Japanese Society for the Promotion of Science (Research Fellowship for Young Scientists, DC1-14J02775 and PD-17J01380, to RY) and from the Natural Sciences and Engineering Research Council of Canada (RGPIN-2016-03711 to SPO). We thank the following people for their helpful comments on previous drafts of the manuscript: Yoh Iwasa, Matt Osmond. Ken Thompson, Sam Flaxman, Mohamad Noor, Bryn Wiley, and three anonymous reviewers. We are grateful to Matt Osmond for suggesting the normalization of AE by BC_L_ as a more informative measure of ecological speciation.

---

**Table 1.** Notation used. Default values are given in square brackets, except where noted in the text.

| Notation | Description |
| --- | --- |
| $m$ | Number of spheres [1] |
| $n$ | Number of traits in each sphere [10] |
| $\mathbf{z}_i = \{z_{i,1}, z_{i,2}, \dots, z_{i,n}\}$ | Phenotype in the $i$th sphere |
| $\mathbf{o}_i = \{o_{i,1}, o_{i,2}, \dots, o_{i,n}\}$ | Optimum phenotype in the $i$th sphere |
| $w(\mathbf{z}_1, \mathbf{z}_2, \dots, \mathbf{z}_m)$ | Individual's relative fitness |
| $q$ | Parameter translating the phenotypic distance from the optimum into the strength of selection [1] |
| $k$ | Degree of epistasis [2] |
| $\mathbf{Z} = \{\mathbf{z}_1, \mathbf{z}_2, \dots, \mathbf{z}_m\}$ | Vector of phenotype sets of all spheres. $\mathbf{Z}_i$ is an initial condition |
| $r$ | Mutation size, drawn from an exponential distribution with mean $\lambda$ |
| $\xi_j$ | Random number drawn from a standard normal distribution |
| $\Delta \mathbf{z}_i = \{\Delta z_{i,1}, \Delta z_{i,2}, \dots, \Delta z_{i,n}\}$ | Phenotypic change due to mutation in the $i$th sphere |
| $\Delta \mathbf{Z} = \{\Delta \mathbf{z}_1, \Delta \mathbf{z}_2, \dots, \Delta \mathbf{z}_m\}$ | Vector of mutational effects across all spheres |
| $s$ | Selection coefficient |
| $h$ | Measure of dominance |
| $N$ | Number of individuals in a population [2000] |
| $c$ | Ploidy level [1 for haploids, 2 for diploids] |
| $u$ | Mutation rate [$10^{-5}$] |
| $b$ | Number of offspring per individual of maximum fitness when calculating establishment probability [5] |
| $\mathbf{O} = \{\mathbf{o}_1, \mathbf{o}_2, \dots, \mathbf{o}_m\}$ | Vector describing the optimal phenotype across all sets of spheres |

**Table S1.**
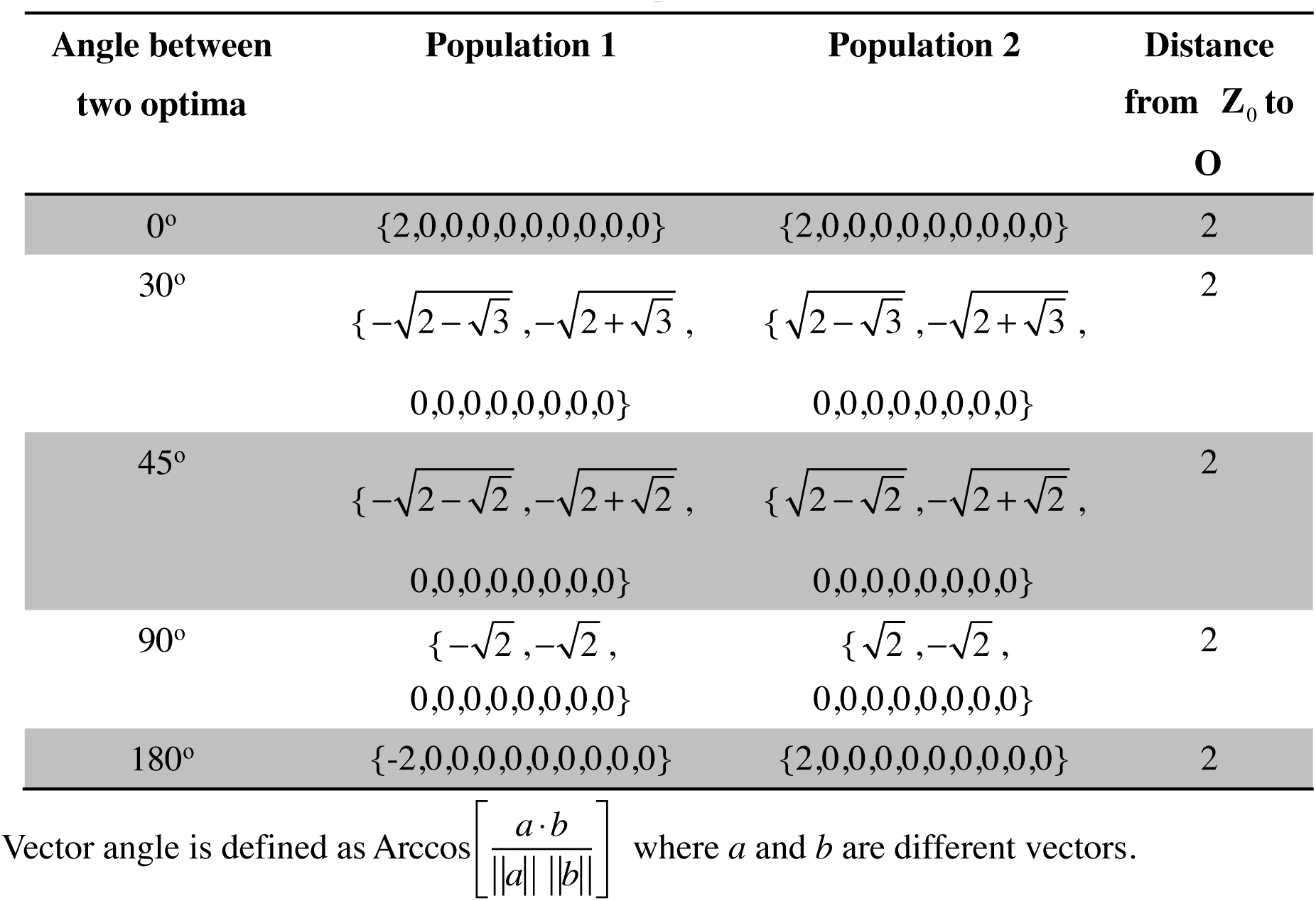
The location of the environmental optima.

**Table S2.**
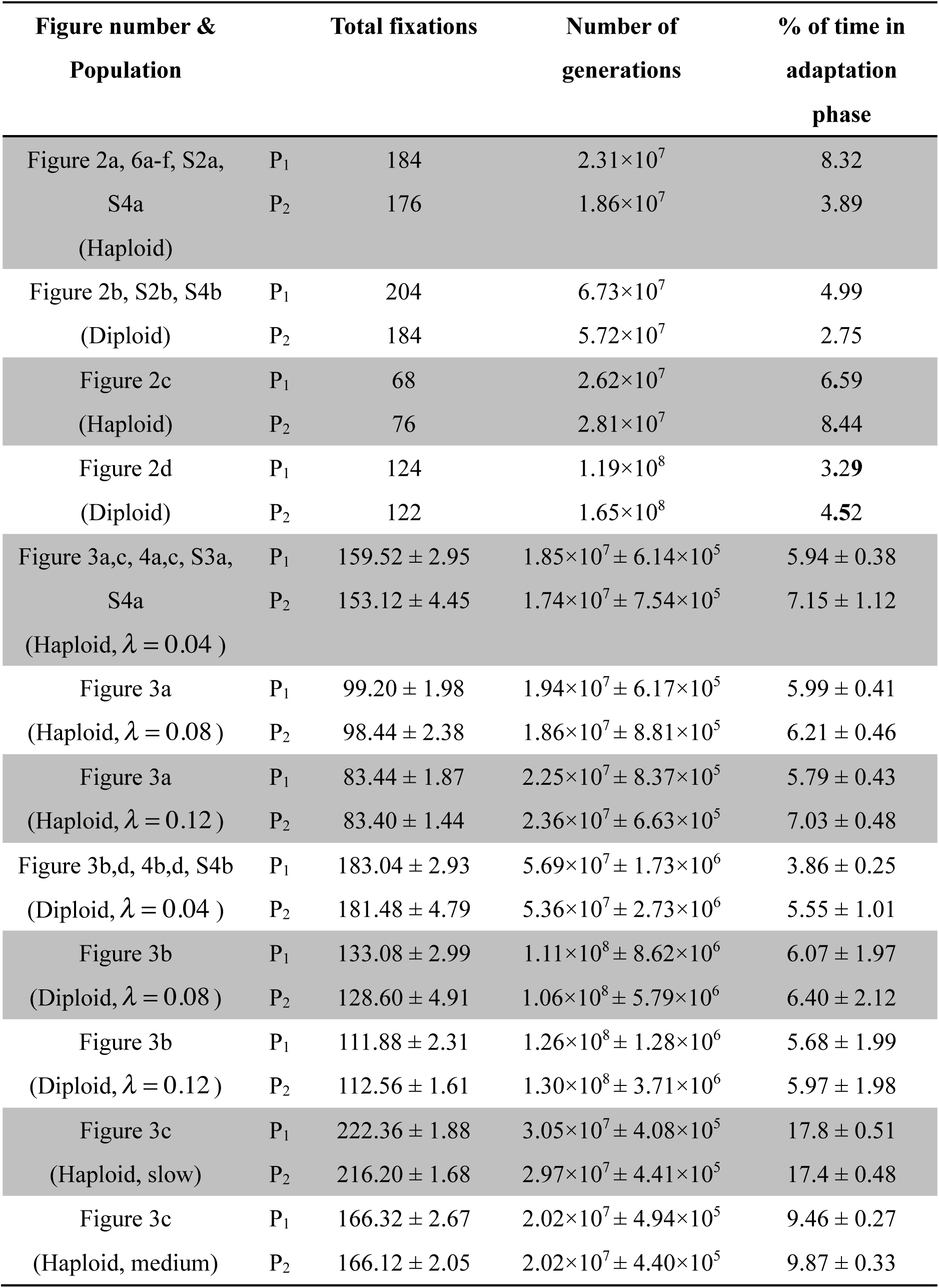
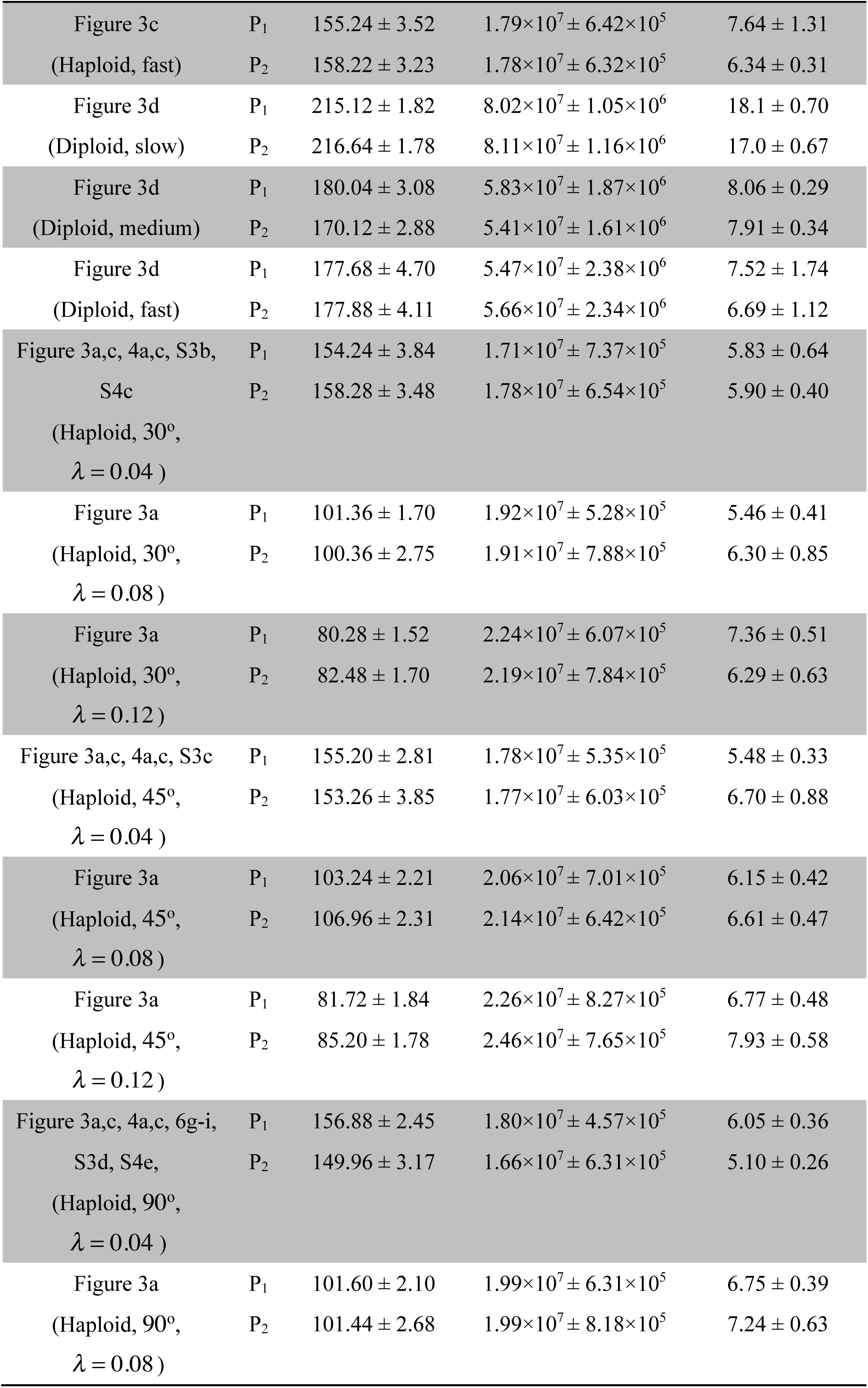
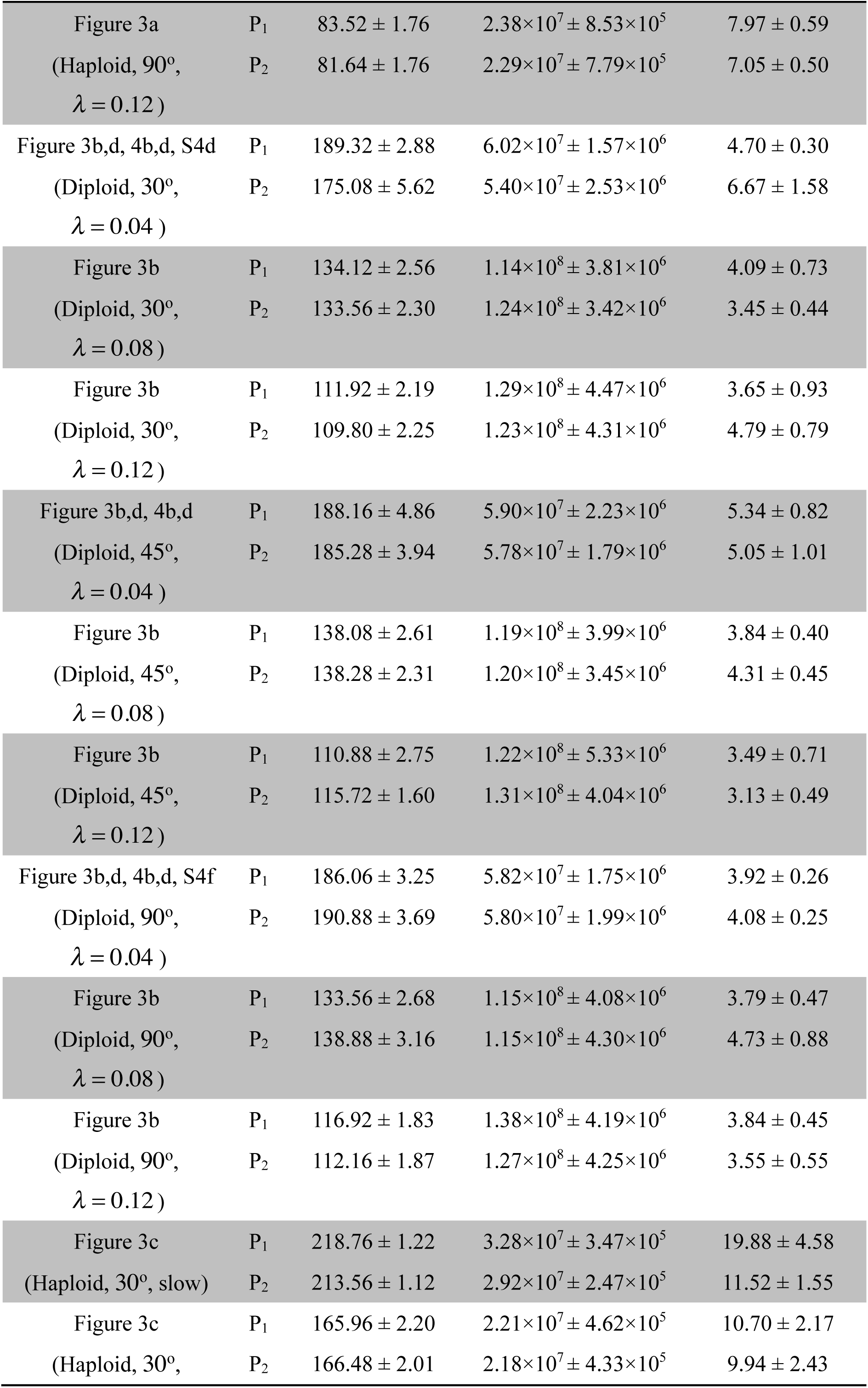
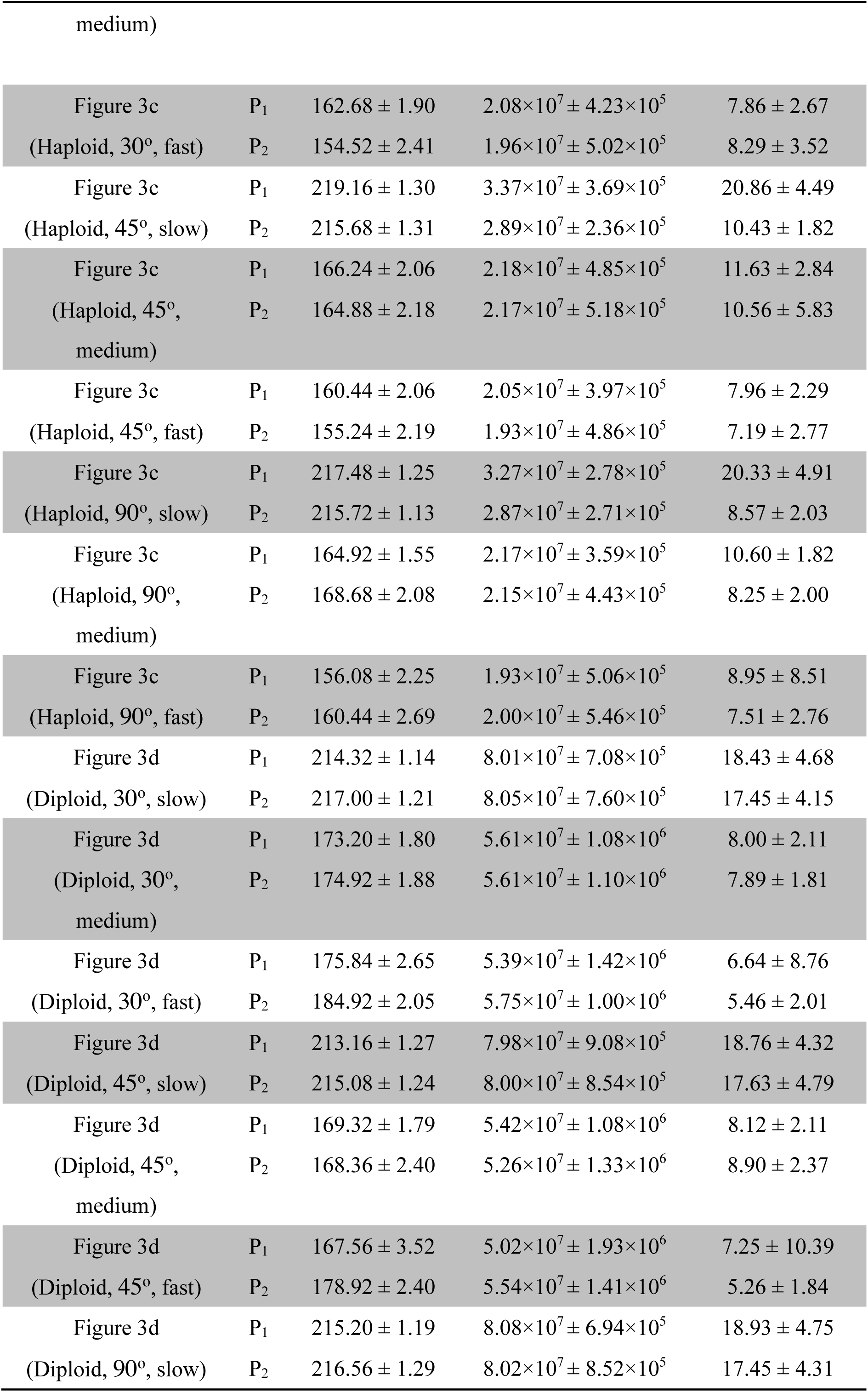
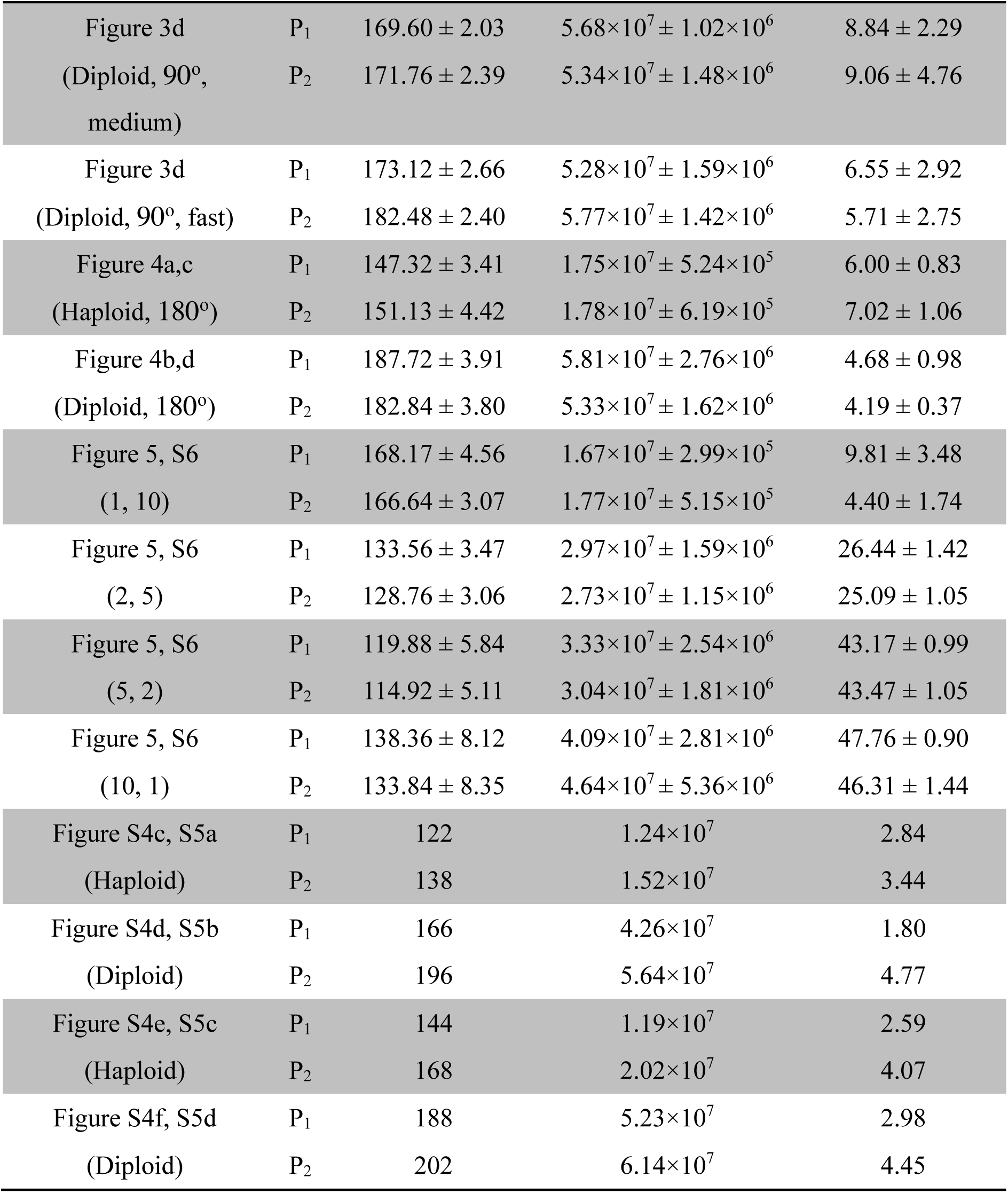
Number of fixation events tracked within the parental populations P_1_ and P_2_ and the corresponding number of generations, based on the expected waiting time until fixation. In each population, we tracked the same total number of fixations events during the adaptation phase and the stationary phase (e.g., 92 for P_1_ and 88 for P_2_ in each phase for the first set of simulations).

**Figure S1.**
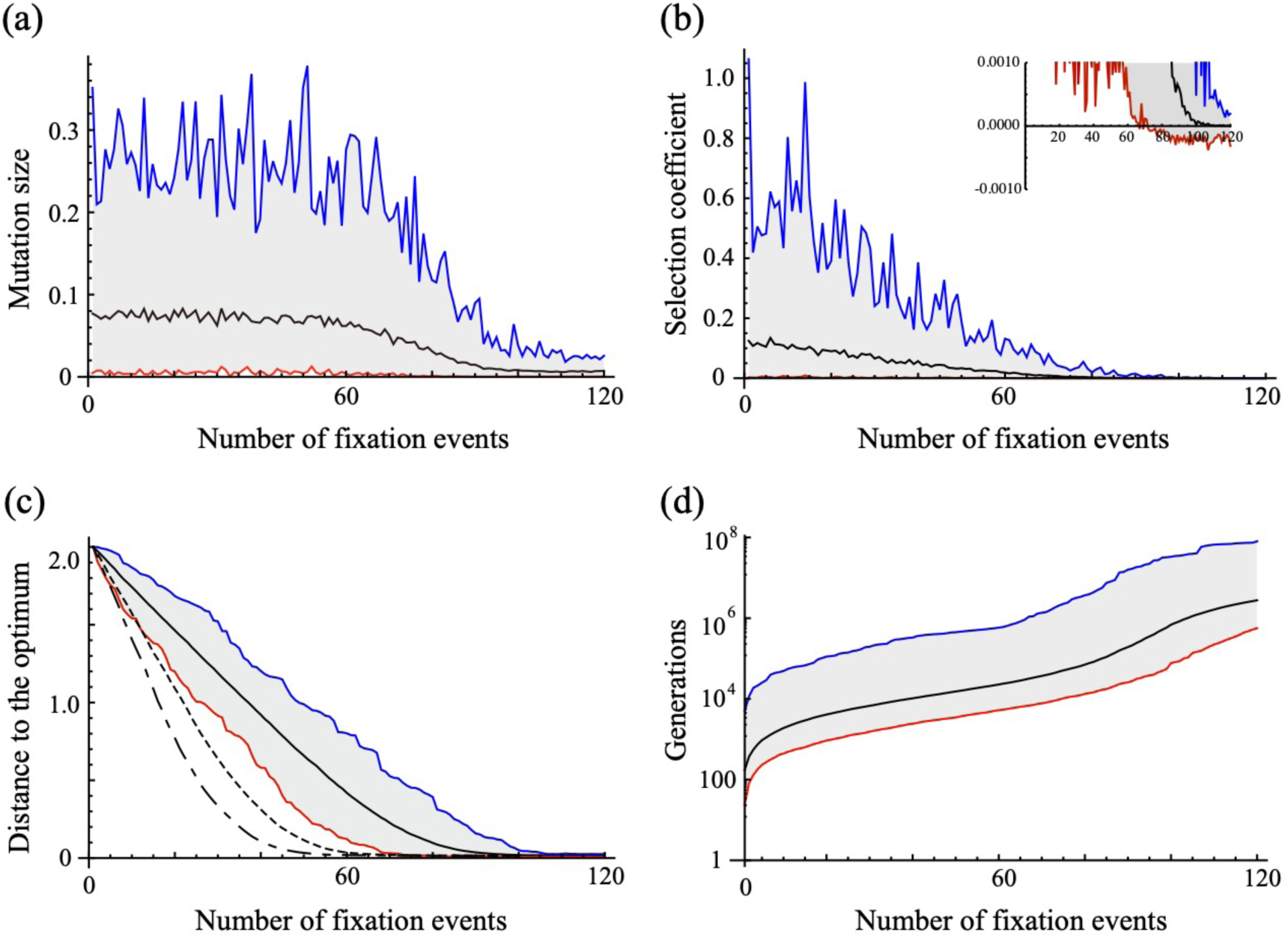
Trajectories within haploid simulations of (a) phenotypic effect size among fixed mutations, (b) selection coefficient, (c) distance to the optimum, and (d) time in units of generations, all plotted against the number of fixation events. Blue, red, and black represent maximum, minimum, and mean, respectively. Selection coefficients are positive in the first “adaptation phase” (taking, on average, 86.9 fixation events) but are more often negative in the second “stationary phase” (inset to b). In (c), mutations are drawn from an exponential distribution with *λ* = 0.04 (right solid line), 0.08 (center dashed line), and 0.12 (left dashed-dotted line). All other panels set *λ* = 0.04. 100 simulations were run for each parameter set. The other parameters are *m*=1, *n*=10, *N*=2000, *q*=1 and *k*=2.

**Figure S2.**
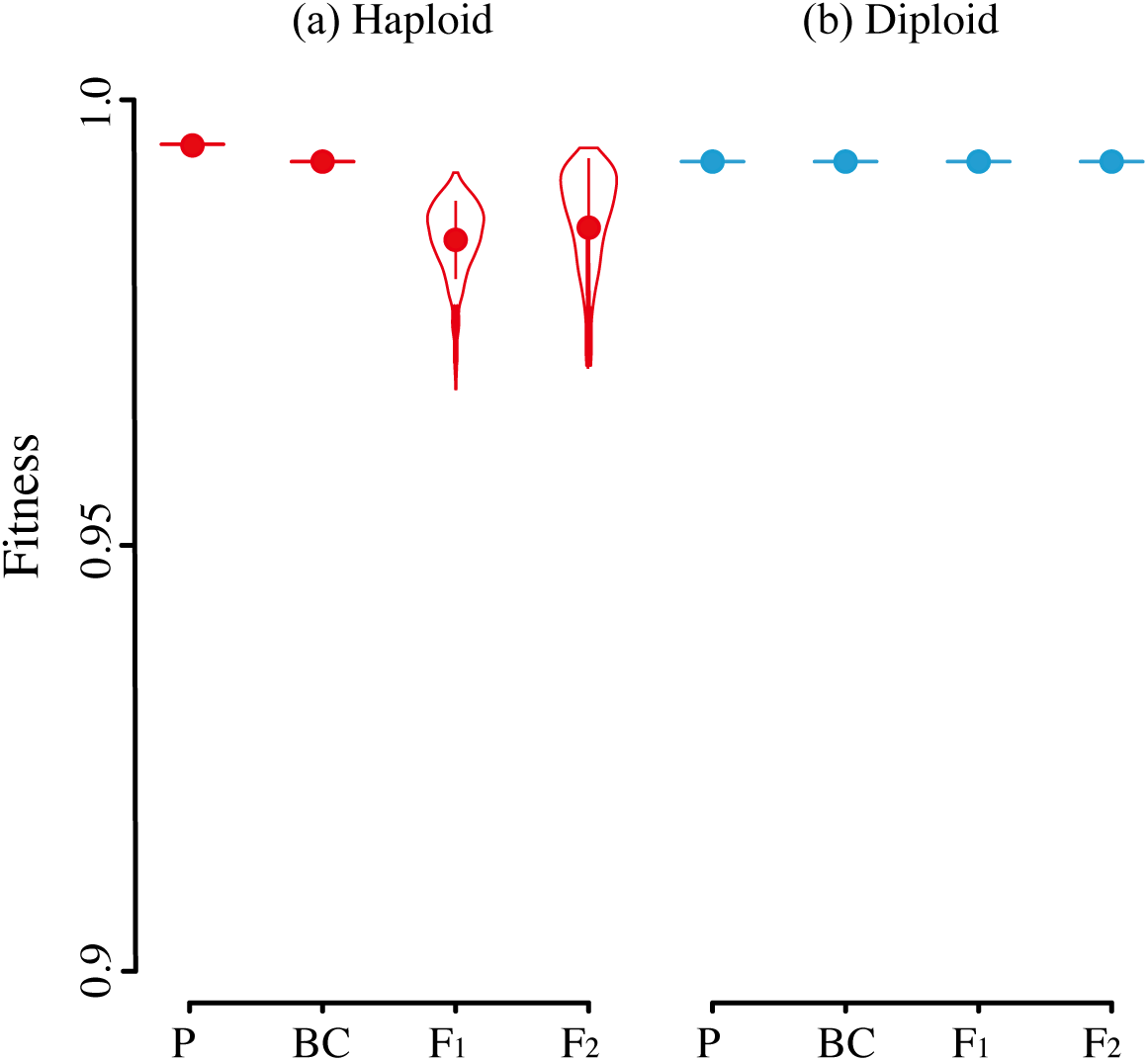
Little reproductive isolation arises during mutation-order speciation during the stationary phase in (a) haploids or (b) diploids. Note that the y-axis range does not start at 0. Data based on the mutations accumulated in Figure 2 (see Table S2 for the number of events). All other parameters are as in Figure 2 with *λ* = 0.04.

**Figure S3.**
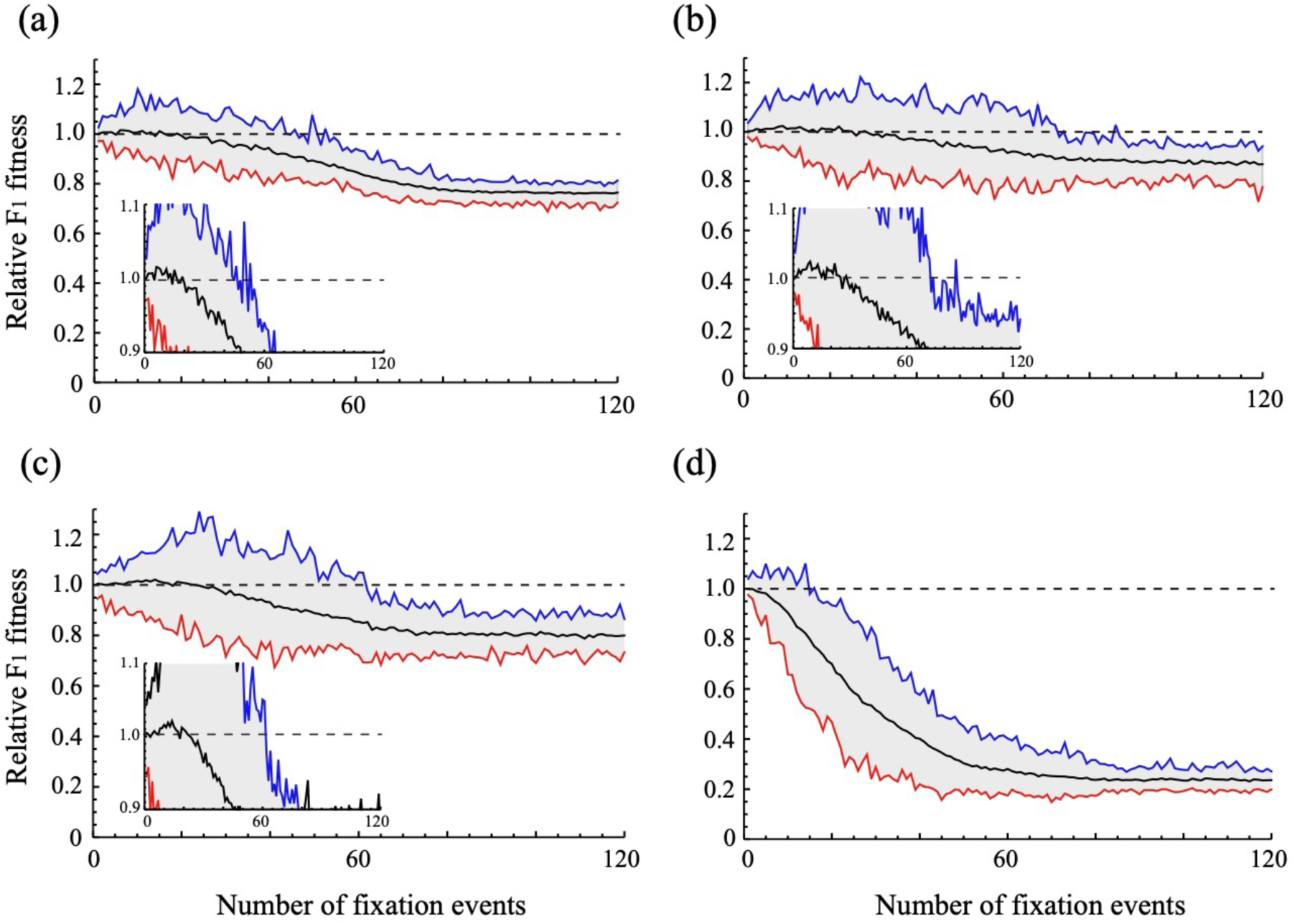
Trajectories of F_1_ fitness relative to the average parental fitness in haploid populations evolving under different environmental scenarios, plotted against the number of fixation events. The trajectories depend on the history of the parental populations: (a) diverged in a constant environment (mutation-order speciation with an angle between the two optima of 0°), (b-d) diverged through adaptation to different environments (ecological speciation with an angle between the two optima of 30°, 45°, and 90°, respectively). Blue, red, and black represent maximum, minimum, and mean, respectively. F_1_ fitness values are shown over time relative to the average fitness of the two parental populations, all measured in patch A at the same time (F_1_ hybrids and parents are equal in fitness, on average, along dashed line at one). Inset figures show the slight heterosis that develops early on. All other parameters are as in Figure S1 except that 50 simulations were run for each parameter set.

**Figure S4.**
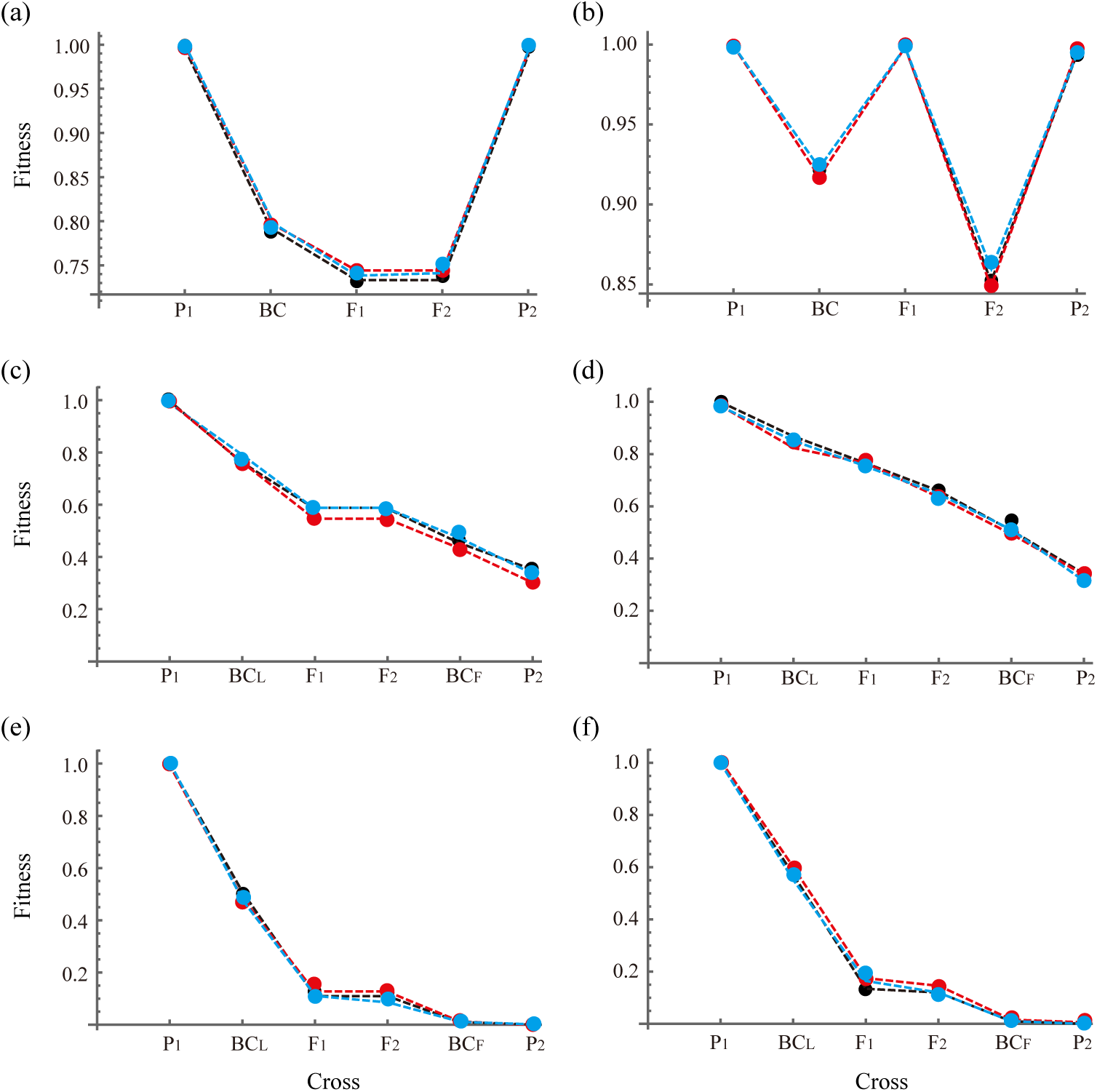
The fitness of hybrids can be predicted using the Brownian bridge approximation of Simon *et al.* (2018). Left and right panel represent haploid and diploid simulations respectively. Top panels show hybrid fitnesses measured in patch A following mutation-order speciation in the same parental environments (as in Figure 2a,b). Remaining panels show hybrid fitnesses measured in patch A following ecological speciation in populations adapting to optima that differ by 30° (middle panels as in Figure S5a,b) or 90 ° (bottom panels; as in Figure S5c,d). The dots represent data generated data in the current paper. The dashed line is predicted by fitting the free parameter E(*S*_†_) of Simon *et al.* (2018) by minimizing the sum of squared deviations between the observed and predicted fitnesses. Three different colors represent three different simulation runs. Choosing E(*S*_†_) based on the class of hybrid with the lowest fitness, as suggested by Simon et al. (2018), does not always provide a good fit, especially when the parents differ in fitness, but fitting the E(*S*_†_) parameter provides a robust predictor to the full set of hybrid fitnesses.

**Figure S5.**
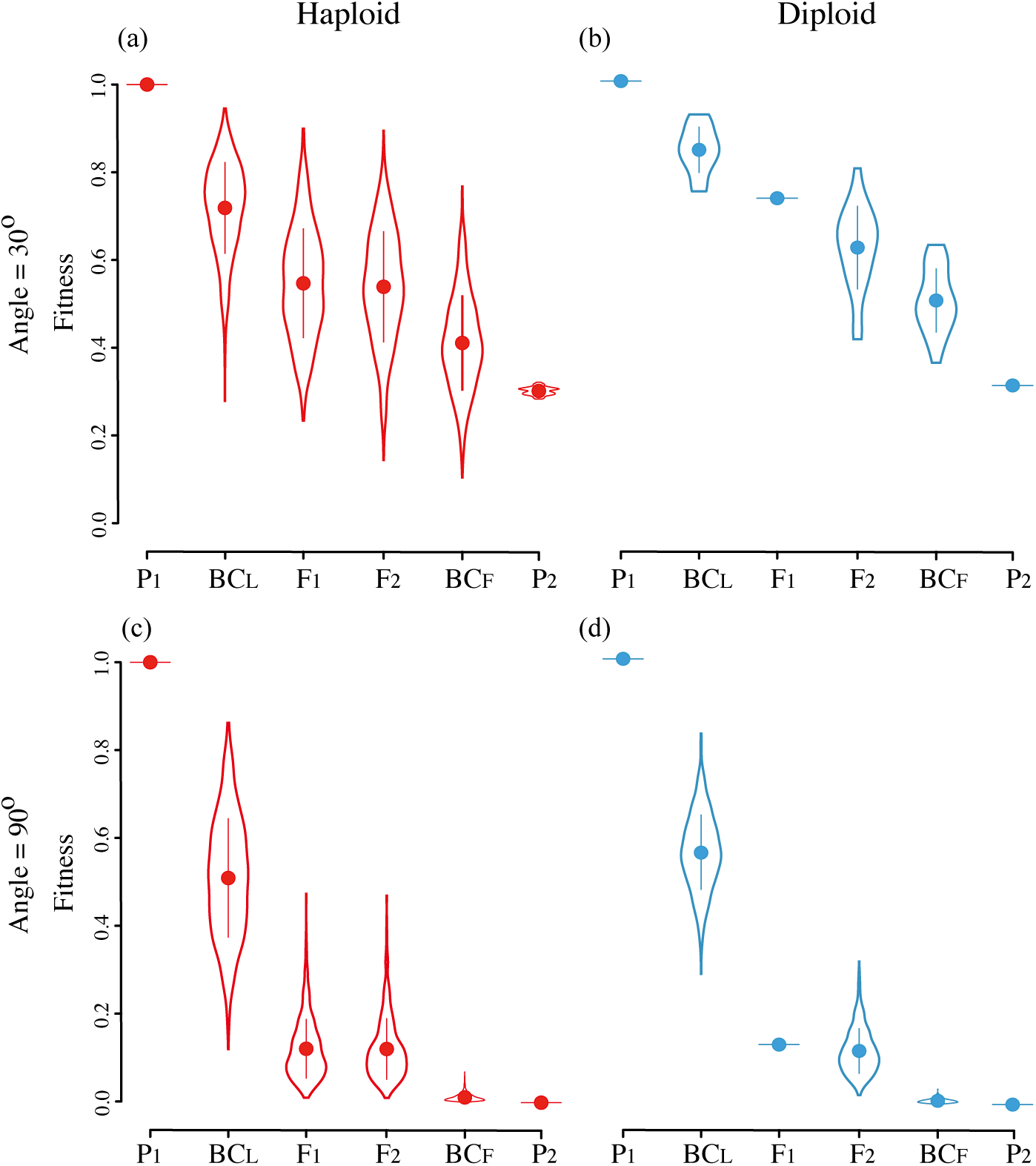
Ecological speciation and the fitness of hybrids in (a, c) haploids and (b, d) diploids. We illustrate the case where the “local” environment is that of population 1 (P_1_), with fitnesses measured in its patch A and using *λ* = 0.04. The angle between the optima of the two environments is set to 30°in top panels and 90°in bottom panels, relative to the origin (see Table S1 for details). All other parameters are as in Figure 2, with distributions and solid points representing the mean ± SD calculated from one simulation. 1000 individuals were generated for each hybrid type.

**Figure S6.**
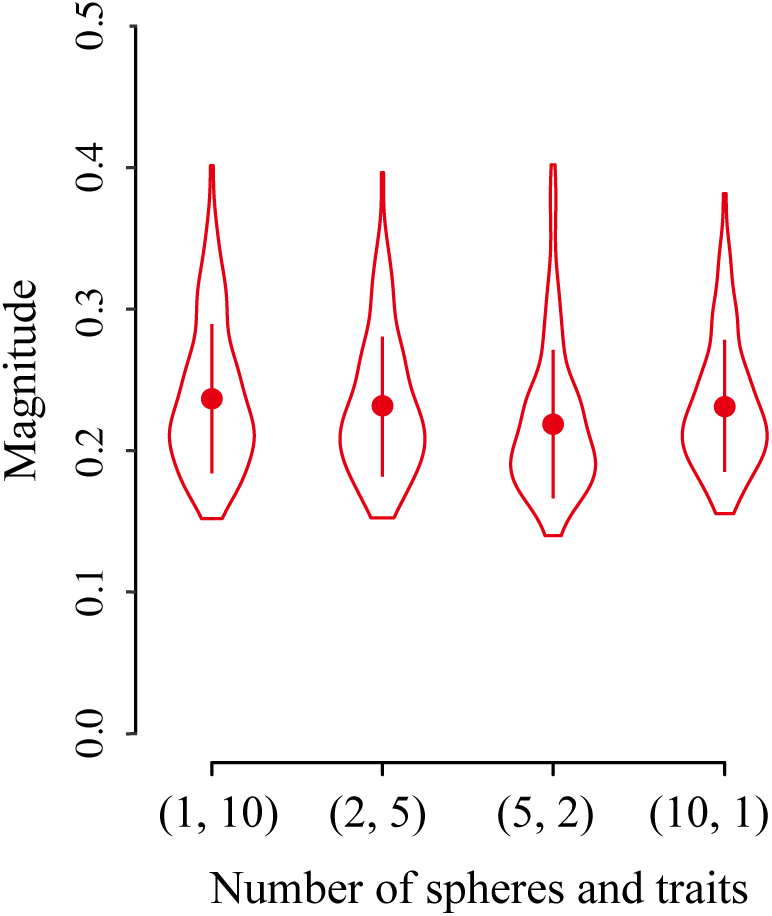
The magnitude of the largest mutation fixed in each of the 50 simulations illustrated in Figure 5 (one-way ANOVA, p>0.05). All parameters are the same as in Figure 5, with distributions and solid points representing the mean ± SD.

